# Kawasemi-Colours: an open end-to-end platform for structure–colour design based on avian structural colour

**DOI:** 10.64898/2026.10.11.758316

**Authors:** Masaki Suimye Morioka

## Abstract

Structural colour provides durable, pigment-free colouration by controlling the interaction of light with nanostructured materials rather than by using wavelength-selective pigments. In birds, non-iridescent blue and green plumage arises from coherent scattering by quasi-ordered keratin–air nanostructures, making these natural systems attractive models for sustainable colour engineering. Despite decades of research on their optical mechanisms, open end-to-end implementations that connect biological nanostructures with colour prediction, inverse design and physics-based evaluation remain limited.

Here I present Kawasemi Colours, an open end-to-end platform for structure–colour design based on avian structural colour. Avian keratin–air nanostructures were formulated as a biologically constrained inverse-design problem and connected to provenance-linked biological data, interpretable forward prediction and physics-based evaluation in an open computational framework. From published electron micrographs, I assembled a provenance-tracked dataset of spongy, channel and quasi-ordered sphere nanostructures with associated colour metadata. I extracted interpretable structural descriptors, developed compact forward models that predict colour from nanostructure images, and implemented an analytic inverse design tool that generates candidate structures for a target colour. Both directions are provided through self-contained interactive web applications.

Kawasemi Colours provides an open framework for studying and designing structural colour from biological nanostructures. By integrating curated data, interpretable machine learning, wave optics, and interactive design tools, the platform lowers the barrier to comparative structural-colour research and provides a cumulative foundation for testing and refining structure–colour relationships as new biological data become available.

## 1. Introduction

Structural colour is a widespread biological phenomenon, occurring across phylogenetically distant groups including insects, birds, fishes, cephalopods and plants (Vukusic and Sambles 2003; Seago et al. 2009; Mäthger et al. 2009; Glover and Whitney 2010; Lloyd and Nadeau 2021). Across these systems, structurally produced colours contribute to diverse biological functions, including camouflage, intra- and interspecific communication and interactions with pollinators, while their repeated evolution provides a tractable system for studying how biological development generates optical function from nanoscale organization (Seago et al. 2009; Mäthger et al. 2009; Glover and Whitney 2010; Lloyd and Nadeau 2021).

Structural colour is generated not by wavelength-selective pigments but by the interaction of light with nanoscale variations in refractive index. Because colour arises from geometry, such systems can be resistant to photobleaching, exhibit high colour purity, and provide naturally evolved design principles for durable, pigment-free optical materials (Vukusic and Sambles 2003; Xiao et al. 2017). Birds provide one of the best-studied examples, particularly the non-iridescent blues and greens produced by quasi-ordered keratin–air nanostructures within feather barbs. Comparative studies across avian lineages have shown substantial diversity in these self-assembled photonic architectures, making them important models for optical physics, evolutionary and developmental biology, and bioinspired materials science (Prum and Torres 2003; Prum et al. 2009; Saranathan et al. 2012).

The optical mechanisms underlying avian structural colour are now well established. Feather barbs contain quasi-ordered β-keratin and air networks that develop through phase separation during feather morphogenesis (Prum et al. 2009; Dufresne et al. 2009). Small-angle X-ray scattering and optical modelling showed that these nanostructures possess a dominant structural correlation length, and that melanin suppresses incoherent scattering and increases colour saturation (Noh et al. 2010; Saranathan et al. 2012; Shawkey and Hill 2006). For the dominant back-scattering peak, reflected wavelength increases with the dominant structural correlation length and depends on the effective refractive index of the keratin–air composite (Noh et al. 2010, eq. 2). Consequently, structural colour can largely be understood as an emergent property of geometry. This relationship also suggests that the colour produced by a nanostructure may be predicted from its geometric characteristics and, conversely, that structural parameters capable of producing a target colour may be inferred from established physical principles.

Despite this well-established physical basis, relatively little work has examined whether these principles can be translated into a generalizable quantitative relationship between naturally occurring nanostructures and their resulting colours, or whether such a relationship can support both forward prediction and inverse structural design. Most previous studies have focused on the description and optical analysis of individual biological structures or on detailed simulations of specific samples. As a result, it remains unclear how far the physical relationships identified in individual systems can be generalized across diverse avian nanostructures and used as practical predictive or design rules.

Over several decades, electron micrographs, structural measurements and colour information have accumulated across studies of diverse biological systems, with individual studies addressing different aspects of the relationship between nanostructure and colour. Bringing these observations together with quantitative optical modelling would provide a basis for systematically comparing how variation in nanoscale organization affects colour. Such comparisons could help investigate whether and how structural changes enable organisms to acquire and diversify structural colours. A unified framework connecting published biological structures, comparative structure–colour analysis and physics-based optical modelling is therefore important for building on this accumulated knowledge and exploring the biological consequences of structural variation.

Here I present Kawasemi Colours, an open platform that converts established principles of avian structural colour into an interpretable framework for colour prediction and structural design. The framework is designed as a cumulative resource: newly reported, provenance-linked structural and optical records can extend its biological coverage, support colour inference for newly characterized feather nanostructures, and provide an evidence base for iterative improvement of predictive models. I curate a provenance-tracked dataset of avian structural-colour electron micrographs together with colour metadata, develop compact forward models that relate nanostructure to colour, implement an analytic inverse design tool for generating candidate structures from target colours, and evaluate the predicted colours using wave optics without fitting colour-specific parameters. The complete workflow is provided through interactive web applications together with openly released code, metadata and reproducible analysis pipelines.

By bringing together curated biological data, interpretable machine learning and first-principles optics, Kawasemi Colours provides a practical route from descriptive biological observations to testable colour predictions and candidate structural designs. I anticipate that this framework will facilitate both biological investigation and the future engineering of self-assembled photonic materials.

## 2. Materials and Methods

### 2.1 Data acquisition and curation

Published electron micrographs of avian structural colour were collected from the PubMed Central (PMC) Open Access subset and bioRxiv. Candidate articles were identified using NCBI E-utilities, followed by manual inspection of figure captions and associated text for electron microscopy (SEM or TEM) of colour-producing feather-barb nanostructures. The curated source set included Parnell et al. (2015), the CC-BY bioRxiv preprint of Bazzano et al. (2020) and its published optical report (Bazzano et al. 2021), Hegyi et al. (2024), Janas et al. (2021), Park et al. (2023), Jeon et al. (2021, 2023) and Sin et al. (2024); record-level provenance is reported in Supplementary Tables S1–S2.

Where a measurable scale bar was present, its pixel length was converted into a pixel-to-micrometer calibration and passed to the extraction of length-valued features. When no reliable scale measurement could be obtained, length-valued image features retain an explicitly labelled heuristic conversion and are not interpreted as absolute measurements; literature-reported structural spacings were retained separately as provenance. Scale-independent descriptors, including structural order, anisotropy and pore fraction, were extracted for all panels irrespective of absolute calibration. Colour-label provenance, peak-wavelength provenance, reported measurement conditions, author-derived versus source values, and scale provenance are tabulated record-by-record in Supplementary Table S2. Display-only sRGB swatches that are not spectrophotometrically calibrated are explicitly flagged and excluded from colour-model training.

Six biological panels include absolute spatial calibration. The remaining panels contribute scale-independent descriptors; their heuristic length conversions and any literature-reported structural spacings are kept distinct from measured image calibration. Complete provenance information is provided in Supplementary Tables S1–S2 and the accompanying metadata tables.

### 2.2 Data licensing and use policy

Image redistribution was assessed separately from information analysis, using image-level credit lines rather than the review article licence alone. Figure 2 reproduces selected avian micrographs under their original CC BY licences. The biomimetic particle panels compiled by Jeon et al. (2021) originate from Kohri et al. (2019), and the Steller’s-jay micrograph originates from Shawkey and Hill (2006). These images, and other third-party panels without documented redistribution permission, are not reproduced here. Their provenance and existing analysis records are retained in Supplementary Tables S1–S2. Restricted images must be obtained from the original sources, subject to applicable permissions; inclusion in an analysis does not grant redistribution rights.

### 2.3 Physics-based structural descriptors

Each grayscale electron micrograph was represented by a seven-dimensional feature vector (FEATS_V1) describing image-derived characteristics of the nanostructure. These descriptors comprise the dominant spatial period, the width of the spatial-frequency distribution (structural order), pore fraction, mean pore diameter, mean wall thickness, anisotropy index and a network-connectivity metric. Length-valued descriptors are expressed in nanometres only when a record-specific scale calibration is available; otherwise their conversion is explicitly heuristic.

The selected descriptors capture the principal structural parameters known to influence coherent scattering in quasi-ordered photonic materials, namely characteristic spacing, structural regularity, porosity and directional organization (Noh et al. 2010; Saranathan et al. 2012). Feature extraction requires no learned parameters and was implemented using OpenCV (Bradski 2000) and NumPy (model/features.py).

### 2.4 Forward prediction models

Two complementary forward models were developed to predict colour from nanostructure images.

The first is a compact convolutional neural network comprising three convolutional blocks followed by a 128-unit fully connected layer (approximately 1.5 × 10^5^ trainable parameters). The network accepts 96 × 96 grayscale image patches and predicts dominant wavelength, saturation and value. Training was performed using synthetic structures generated from physically parameterized quasi-ordered textures, followed by fine-tuning using the biological dataset. Computation was accelerated using the Apple Silicon Metal Performance Shaders (MPS) backend where available.

The second model is an interpretable ridge-regression framework operating on the FEATS_V1 descriptor vector. Unlike the CNN, this approach provides a direct relationship between individual structural parameters and predicted colour, facilitating physical interpretation.

Synthetic training structures were generated on the fly by superposing plane waves with prescribed spatial frequencies and orientation distributions, followed by thresholding to achieve the desired pore fraction (model/synth.py). Each run used 800 in-memory 96 × 96-pixel structures generated with seed 0. Pore fraction was sampled uniformly from 0.30– 0.70, disorder from 0.05–0.75, anisotropy from 0–0.6, and a visible-band target peak was converted to a characteristic spacing clipped to 90–320 nm. These 800 generated structures are distinct from the 11 synthetic-proxy metadata records in the curated dataset. This procedure produces spongy nanostructures with known generator parameters.

Model performance was evaluated with tests matched to the question addressed by each approach. For the interpretable regressor, two random five-fold analyses quantified performance with different evaluation populations: train_real.py evaluates 39 eligible records (28 real images and 11 synthetic-proxy metadata records), whereas validate_real_grouped.py evaluates the 28 eligible real images. In both analyses, 800 on-the-fly synthetic structures and the non-held-out curated records are available for training. The real-image analysis was also repeated with leave-one-species-out (LOSO) and leave-one-paper-out (LOPO) validation to test transfer to an unobserved recorded species or source paper. Random folds use np.random.default_rng(0) followed by five contiguous array_split folds, and the real-image membership is archived in Supplementary Table S4. For the convolutional neural network, evaluation on held-out synthetic structures after fine-tuning on the biological dataset quantifies image-based prediction within the synthetic generator domain.

### 2.5 Inverse structural design

An analytical inverse-design framework was implemented to estimate candidate nanostructures from a target colour. Previous work used machine learning to map target colours to engineered dielectric-array geometries (Huang et al. 2019); here, the inverse problem was restricted to biologically informed keratin–air candidates within the declared generator priors. The target colour was first converted into dominant wavelength, saturation and value, and an initial characteristic spacing was then calculated using the first-order dominant-backscattering approximation used for quasi-ordered avian photonic nanostructures (Noh et al. 2010, eq. 2):

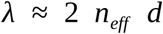

where *d* is the characteristic structural spacing and *n_eff_* is the effective refractive index of the keratin –air composite.

The remaining structural parameters, including pore fraction and structural order, were subsequently refined by optimizing agreement between the analytical forward model and the target colour. Because colour-only inversion is non-unique, the output is one prior-constrained candidate structure. The reported spacing range describes model-conditioned variation under the declared pore-fraction prior (0.30–0.70) through 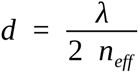.

The resulting parameters are exported as lightweight JSON files and visualized through an interactive web application. Users can specify an arbitrary target colour and obtain the predicted nanostructure together with its geometric cross-section, reflectance spectrum and reconstructed colour in real time.

### 2.6 Physics-based colour computation

Colour was computed directly from nanostructure geometry, independently of the machine-learning models, using the coherent single-scattering framework developed for quasi-ordered biological photonic structures (Prum and Torres 2003; Noh et al. 2010).

Electron micrographs or synthetic structures were interpreted as refractive-index distributions composed of keratin (n = 1.56) and air (n = 1.00). The radially averaged Fourier power spectrum of each structure was calculated to obtain the structure factor, S(q), which characterizes the dominant spatial frequencies within the quasi-ordered material.

Using a dominant-backscattering approximation to the reciprocal-space single-scattering framework of Noh et al. (2010, eqs 2–3), the implementation estimates the relative reflectance spectrum as

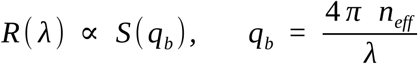

The dominant reflected wavelength was calculated using the first-order relation given above.

The 25th percentile of positive sampled spectral values was subtracted and negative values were clipped to zero before normalization. This baseline correction is a heuristic adopted in the present implementation, not a quantitative model of melanin absorption. Reflectance spectra were converted into CIE XYZ tristimulus values using the CIE 1931 2° standard observer with the analytical approximation of Wyman et al. (2013), followed by conversion into sRGB under a D65 illuminant.

No parameters were fitted to the displayed target colours. The forward calculations instead use declared literature-informed spacings, representative pore fraction, preprocessing, and generator priors; the same optical model is used throughout the forward simulations, inverse design framework, and interactive web applications.

The colour visualizations used the CIE-derived sRGB value from each simulated reflectance spectrum. For colourized structure images, hue and saturation were taken from the simulated sRGB value while the grayscale intensity of the synthetic structure or electron micrograph was retained as the value channel. Uniform colour patches were generated directly from the same simulated sRGB value without added texture, shading or highlights. Record-associated sRGB annotations were displayed separately as inset squares.

For an idealized ordered multilayer, angle-dependent reflectance was calculated with the standard 2 × 2 characteristic-matrix transfer-matrix method for stratified media (Yeh 1988). The model used alternating refractive indices of 2.0 and 1.40, a thickness of 70 nm for each layer, eight high/low pairs (16 layers total), an incident medium with n = 1.0 and a substrate with n = 1.5. Reflectance was evaluated from 360 to 780 nm in 2 nm steps at incidence angles of 0°, 15°, 30°, 45° and 60° and reported as the unpolarized mean of the s- and p-polarized responses. This ordered-stack model was used to characterize the angle-dependent spectral response of an iridescent mechanism, distinct from the quasi-ordered keratin–air single-scattering calculations.

### 2.7 Three-dimensional structural analysis

Three-dimensional reconstruction and Fourier-based colour prediction have previously been demonstrated for an avian spongy keratin–air nanostructure (Shawkey et al. 2009). Here, rather than reconstructing a measured volume, synthetic isotropic keratin–air structures were generated as cubic binary voxel volumes. Each volume used for the optical calculations contained 72 × 72 × 72 voxels at 20 nm per voxel, corresponding to a cube 1.44 µm on each side. The pore fraction was fixed at 0.50, the disorder parameter at 0.12, and the random seed at 1. Volumes were generated for characteristic spacings *d* = 160, 200 and 250 nm.

Each volume was generated from the sum of 40 three-dimensional cosine waves. For voxel coordinate x = (x, y, z), the continuous field was defined as

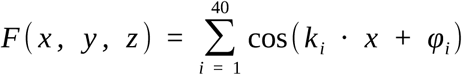

The propagation direction and phase of each wave were sampled independently at random. Its spatial frequency was centred on 1/*d* and varied uniformly by ±0.9 times the disorder parameter (±10.8% under the setting used here). The summed field was centred and scaled to unit variance, then thresholded at the quantile defined by the pore fraction. Voxels at or above the threshold were assigned to keratin (1), and voxels below the threshold to air (0), producing an interpenetrating quasi-ordered three-dimensional network.

For each 723 volume, the mean voxel value was subtracted and a three-dimensional Hann window was applied before computing the three-dimensional fast Fourier transform. The squared Fourier magnitude was radially averaged to obtain the structure factor S(q). Reflectance was sampled using the back-scattering relation defined in Section 2.6 and normalised, with *n_eff_* = 1.28 for the 0.50 pore fraction. The resulting spectra were converted to CIE-based sRGB values using the same workflow as the two-dimensional analytical calculations.

The Figure 8 mid-plane slices and reflectance spectra were obtained from the 723 volumes used for the optical calculations. The upper-row surface views were rendered from keratin boundary voxels in separate 403 volumes generated with the same *d*, pore fraction, disorder and random seed.

### 2.8 Qualitative FDTD cross-check of the analytical model

Finite-difference time-domain (FDTD) simulations were performed using MEEP (version 1.34.0; Oskooi et al. 2010). Three-dimensional keratin–air voxel models were converted into MaterialGrid representations for the full-wave electromagnetic calculations.

Each model was illuminated with a broadband plane wave. Reflectance spectra were calculated from the ratio of reflected to incident flux and converted into colour using the same CIE-based workflow as the analytical calculations. The 243-voxel simulation cell was periodically repeated in x and y and terminated by perfectly matched layers along z.

Numerical sensitivity was assessed for the *d* = 185 nm structure at resolutions of 24, 36, and 50 pixels/µm, with and without subpixel smoothing. Multiple structure realisations were also evaluated to assess seed dependence. The qualitative comparison examined the spectral region and spacing-dependent trend of the simulated reflectance peaks relative to the analytical predictions. Full commands, software versions, boundary conditions, voxel size, resolution settings, and per-seed outputs are reported in Supplementary S5.

## 3. Results

### 3.1 Overview of the study and open outputs

The study was organized as a connected evidence chain from source data to design output, with five stages: curation of provenance-linked electron micrographs and colour metadata, extraction of structural descriptors, prediction of colour from real images, physics-based tests of the spacing –wavelength relation, and delivery through forward and inverse design tools (Figure 1). Each stage addresses the question needed for the next. Grouped cross-validation tests prediction on real-image records; synthetic spacing sweeps test whether controlled geometric changes produce the expected spectral trend; explicit three-dimensional volumes and qualitative MEEP finite-difference time-domain (FDTD) calculations cross-check that trend with more detailed optical models; and transfer-matrix calculations determine how an ordered multilayer responds to viewing geometry.

**Figure 1.**
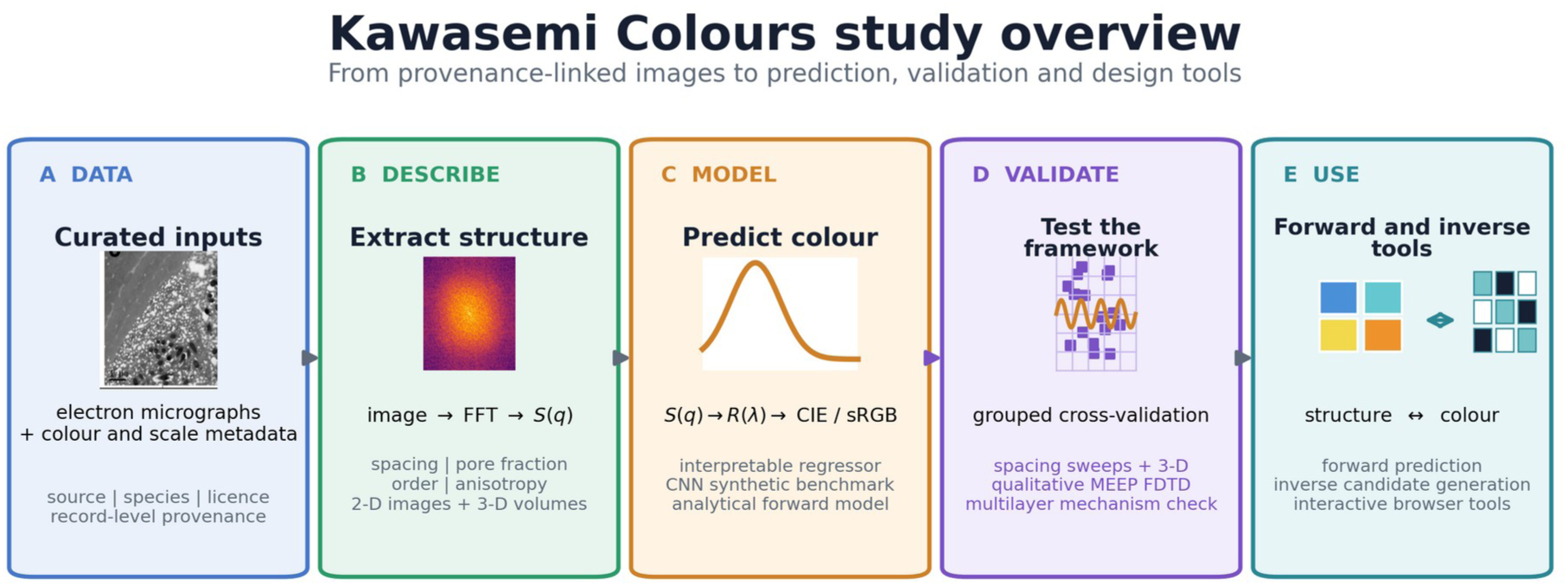
Study overview. (A) Provenance-linked electron micrographs and associated colour and scale metadata form the curated inputs. (B) Structural descriptors are extracted from two-dimensional images and three-dimensional volumes. (C) Structure-factor and statistical models predict reflectance and colour. (D) Grouped cross-validation, synthetic spacing sweeps, three-dimensional calculations, qualitative MEEP FDTD and a multilayer calculation evaluate complementary aspects of the framework. (E) Forward and inverse browser tools link structure with predicted colour and generate candidate structures from target colours. The Eurasian-jay micrograph is reproduced from Parnell et al. (2015; CC BY 4.0).

The GitHub repository contains data-curation and feature-extraction code, model-training and grouped-validation scripts, analytical, three-dimensional and MEEP workflows, figure-generation code, and two self-contained browser tools for structure-to-colour and colour-to-candidate-structure exploration. The versioned software and associated data archive is available on Zenodo (doi:10.5281/zenodo.23445928). The image-level rights assessment is reported in Supplementary Tables S1–S2; third-party images retain their original terms and must not be treated as covered by the software licence or a blanket repository licence.

### 3.2 A curated dataset of avian structural colour

The biological and comparison populations used in the predictive analyses were defined by a provenance-tracked dataset of 42 structural-colour records: 31 real-image records (26 avian micrographs and five biomimetic comparison images) and 11 synthetic-proxy metadata records used for model development. The avian records include 14 named species and four panels recorded only at the avian structural-type level (Figure 2). The primary physical analysis concerns non-iridescent keratin–air morphologies—spongy, channel and quasi-ordered sphere nanostructures.

**Figure 2.**
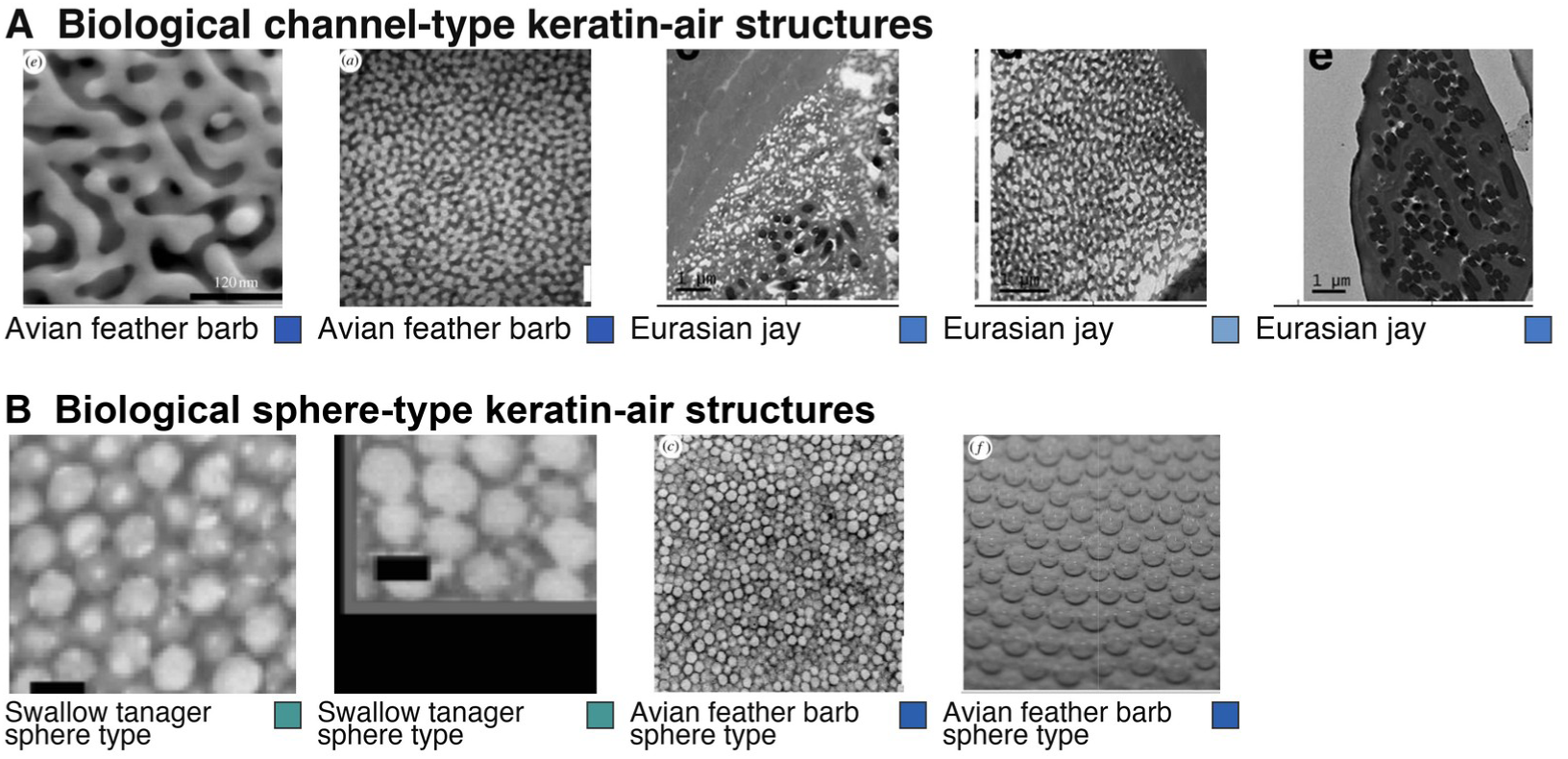
Representative avian micrographs from the curated dataset. (A) Biological channel-type keratin–air structures. (B) Biological sphere-type keratin–air structures. Each record links an electron micrograph to structural classification, colour information and source provenance. Coloured squares show record-associated sRGB values and are not a quantitative scale. Panel A, left to right: Prum et al. (2009; two images, CC BY 2.0) and Parnell et al. (2015; three images, CC BY 4.0). Panel B, left to right: Bazzano et al. (2020; two images, bioRxiv preprint, CC BY) and Prum et al. (2009; two images, CC BY 2.0). The selected images have been cropped and arranged; source scale bars are retained where present. Complete record-level provenance and redistribution conditions are provided in Supplementary Tables S1–S2.

A melanosome-array group was included to examine whether a model built around keratin–air morphology transfers to a distinct optical mechanism. These ordered melanin-rich multilayers in iridescent feather barbules produce colour mainly by multilayer interference and therefore serve as an out-of-domain comparison. Four common-pheasant records were extracted from Jeon et al. (2023), and five records for rock dove, Lawes’ parotia, green-winged teal, violet-backed starling and white-booted racket-tail hummingbird were extracted from figures compiled in Jeon et al. (2021). The latter originated from Yoshioka et al. (2007), Stavenga et al. (2011a), Xiao et al. (2017), and Eliason et al. (2013, 2020); record-level figure provenance is given in Supplementary Table S2.

Each record was structured to make the subsequent comparisons traceable, with fields for structural class, colour metadata, publication source, licence information and, where available, absolute spatial calibration. An illustrative sRGB display swatch and a separately sourced dominant reflectance peak are retained as distinct fields rather than treated as mutually derivable. Display swatches without quantitative calibration are flagged and excluded from colour-model training. Six biological panels provide directly measurable scale calibration; the other panels contribute scale-independent descriptors, with heuristic image lengths and literature-reported spacings explicitly identified in the provenance table.

This curation produced a provenance-tracked resource that combines avian nanostructure, colour and image-level licensing information in one analysis-ready framework. The metadata distinguish the full analysis population from images eligible for redistribution; the prediction analyses below retain the original evaluation populations.

### 3.3 Predicting colour from biological nanostructures

The same forward model and its inverse counterpart are exposed through the two browser tools shown in Figure 3. Prediction from real images was evaluated by asking whether interpretable features extracted from electron micrographs recover the associated peak wavelength. The feature-based ridge regressor was tested on 28 real-image records eligible for colour-model training (Figure 4). The population comprised 14 mechanism-matched non-iridescent keratin–air records, nine melanosome-array records forming the out-of-domain mechanistic comparison group, and five biomimetic comparison records; three records retaining only illustrative, non-calibrated sRGB swatches were excluded. Scale strata, fold assignments, seeds and generator settings are reported in Supplementary Tables S3 and S4.

**Figure 3.**
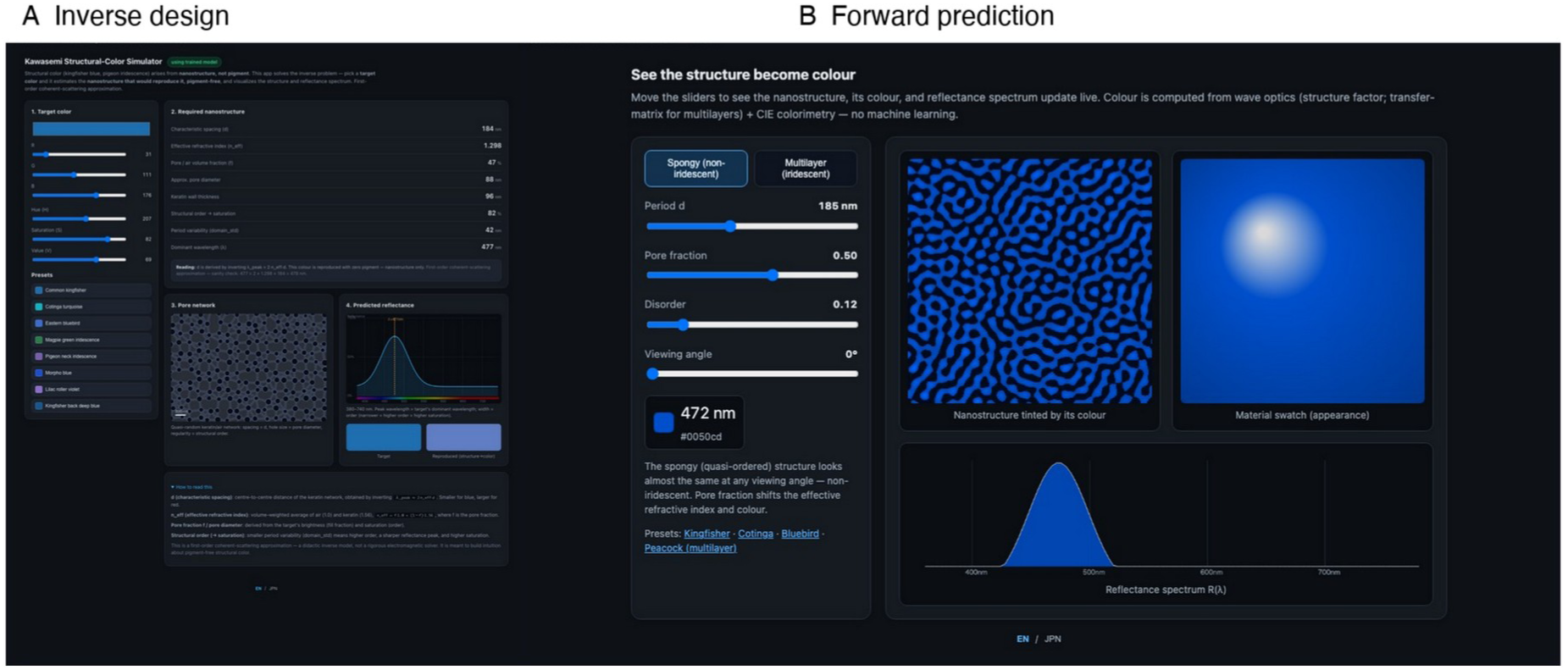
Representative interfaces of the Kawasemi Colours browser tools. (A) The inverse-design interface converts a target sRGB colour into candidate nanostructural parameters, a generated structure and its forward-modelled reflectance. (B) The forward interface converts structural parameters into a nanostructure, reflectance spectrum, CIE-derived colour and rendered material appearance. The displayed spectra, colours and appearances are model outputs.

**Figure 4.**
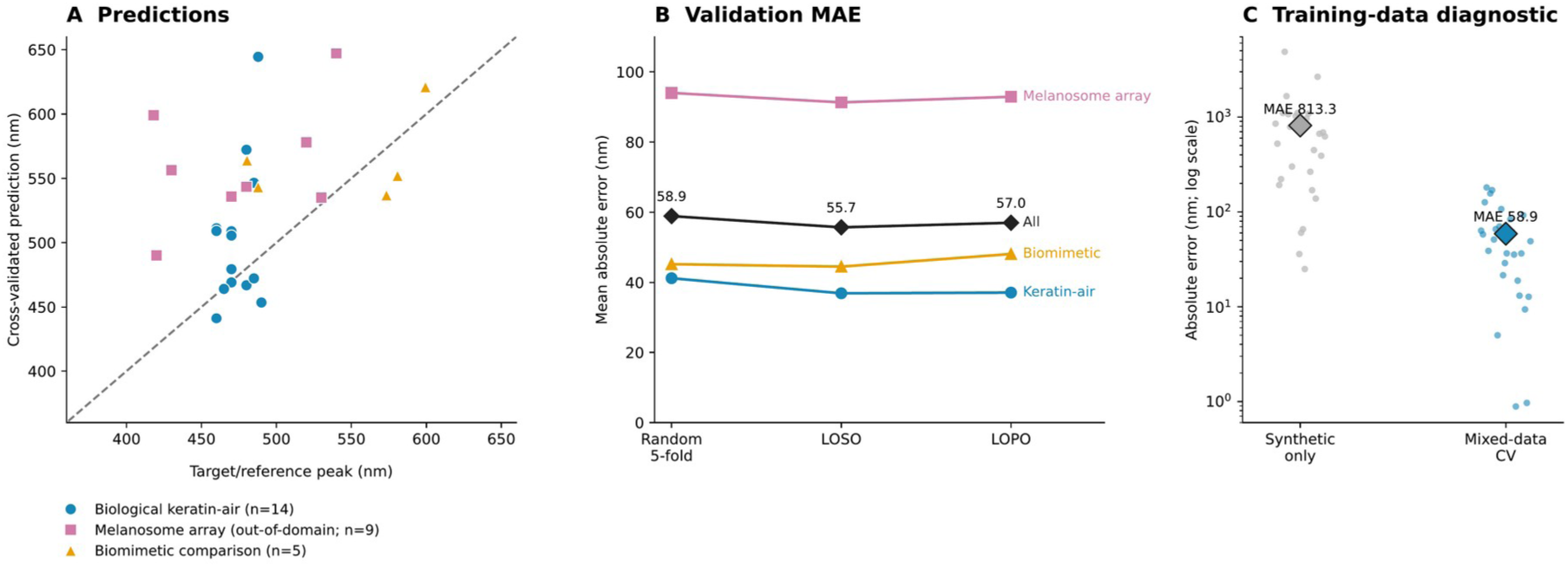
Cross-validated peak-wavelength prediction and training-data comparison. (A) Target/reference peak wavelength versus prediction for the archived seed-0 random five-fold analysis of 28 eligible real-image records; the dashed line is the identity line. Symbols distinguish 14 biological non-iridescent keratin-air records, nine melanosome-array records analysed as an out-of-domain mechanistic comparison (Jeon et al. 2021, 2023), and five biomimetic records. (B) Mean absolute error for the same records under random five-fold, leave-one-species-out (LOSO), and leave-one-paper-out (LOPO) validation. The black series is pooled across records; coloured subgroup series describe strata from the same pooled fits. (C) Absolute errors for the synthetic-only ridge model and mixed-data random five-fold model evaluated on the same 28 records; diamonds show means and the ordinate is logarithmic. Nineteen of 28 unconstrained synthetic-only predictions fell outside the 380–750 nm visible range.

Generalization across related records was assessed by comparing random five-fold, leave-one-species-out (LOSO) and leave-one-paper-out (LOPO) validation. Across the 28 real images, random five-fold validation yielded a mean absolute error (MAE) of 58.9 nm and a median absolute error of 50.1 nm. LOSO and LOPO validation yielded MAEs of 55.7 nm (median 40.1 nm) and 57.0 nm (median 41.1 nm), respectively (Figure 4A,B). The similar MAEs across the three schemes show that holding out the recorded species or source paper did not increase the average error relative to random folds.

Prediction accuracy differed across structural mechanisms within the same pooled-model predictions. The biological keratin–air subset had MAEs of 41.2, 36.9 and 37.1 nm under random five-fold, LOSO and LOPO validation, whereas the melanosome-array comparison group had corresponding MAEs of 94.0, 91.3 and 92.9 nm; the biomimetic comparison group yielded 45.2, 44.5 and 48.1 nm (Figure 4B). Thus, the feature model was most accurate for the keratin–air mechanism it was designed to describe and transferred less accurately to multilayer melanosome arrays. In a complementary random five-fold evaluation of all 39 eligible records, including 11 synthetic-proxy records, the MAE was 57.2 nm and the median absolute error was 37.0 nm (Supplementary Table S4).

Synthetic-to-real transfer was then tested by applying the synthetic-only ridge model to the same 28 real-image records. Its raw MAE was 813.3 nm, and 19 of 28 predictions fell outside the 380– 750 nm visible range, whereas mixed-data random five-fold validation yielded an MAE of 58.9 nm (Figure 4C). Leaving the synthetic-only outputs unconstrained exposed the extrapolation directly and showed that incorporating biological records was necessary to bridge the synthetic-to-real domain shift in this model.

Image-based CNN performance within the synthetic population was quantified using 120 seed-0 structures held out from fitting. After biological fine-tuning, the checkpoint achieved an MAE of 65.0 nm and a median absolute error of 63.5 nm. These values provide the synthetic-population performance estimate for the fine-tuned CNN.

With the forward prediction behaviour characterized, the inverse tool was used to ask which feather-like geometry could produce a specified colour. For a kingfisher-like blue centred at 467 nm, the model returned a prior-constrained candidate spacing of approximately 183 nm at the representative pore fraction, together with a wider model-conditional range when pore fraction was varied within its declared prior. Evaluation with the same forward physical model reproduced the target colour, motivating the controlled spacing tests in the next section.

### 3.4 Characteristic spacing shifted simulated reflectance peaks

Characteristic spacing was then varied in a two-dimensional synthetic sweep to test the expected spectral trend independently of image-level regression, while the remaining generator settings were held fixed. Increasing *d* from 140 to 270 nm shifted the ensemble-averaged reflectance peak from 394 to 756 nm (Figure 5A). The five simulated peaks were 394, 410, 492, 614 and 756 nm at *d* = 140, 165, 190, 225 and 270 nm, respectively. The corresponding first-order estimates from *λ ≈* 2 *n_eff_ d* were 358, 422, 486, 576 and 691 nm, giving absolute differences of 6–65 nm.

**Figure 5.**
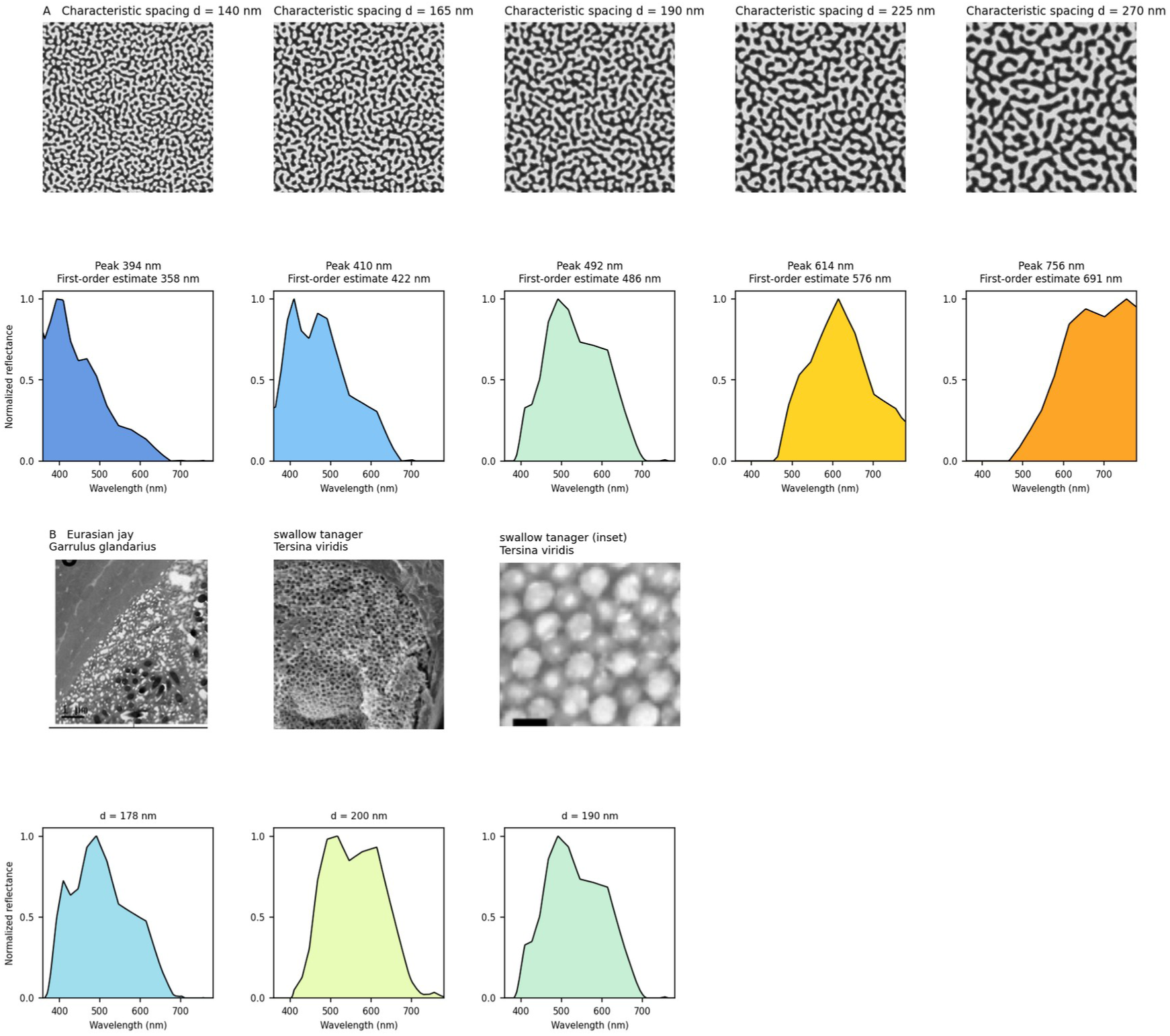
Physics-based in-silico consistency of structural colour. (A) Synthetic keratin-air structures with increasing characteristic spacing show systematic shifts in the simulated reflectance peak. The top row shows one representative realisation at each spacing; the spectra are normalized ensemble means across six independently generated structures. Labels compare the peak of each ensemble-averaged spectrum with the corresponding first-order estimate. (B) Published micrographs establish species and panel context for literature-informed synthetic candidates. The lower panels show normalized ensemble-mean spectra calculated for these candidates using the stated spacing and pore-fraction parameters. For *d* = 178, 200 and 190 nm, respectively, the simulated peaks were 492, 516 and 492 nm, the literature-derived target peaks were 480, 550 and 550 nm, and the absolute differences were 12, 34 and 58 nm. The Eurasian-jay and swallow-tanager micrographs are reproduced from Parnell et al. (2015; CC BY 4.0) and Bazzano et al. (2020; bioRxiv preprint, CC BY), respectively.

The same spacing-to-spectrum calculation was next applied to three record-level spacing and peak-wavelength pairs to examine agreement with literature-linked colour targets (Figure 5B). Candidate structures with *d* = 178, 200 and 190 nm produced ensemble peaks of 492, 516 and 492 nm, compared with literature-derived target peaks of 480, 550 and 550 nm, giving absolute differences of 12, 34 and 58 nm. The micrographs in Figure 5B identify the biological sources of those candidate parameters, so the panel connects each quantitative spacing–peak test to the corresponding feather nanostructure.

Generator-driven variability was measured by repeating each candidate 20 times at the same nominal parameters. The three record-linked settings yielded ensemble peaks of 468, 518 and 492 nm, with between-realisation standard deviations of 32.6, 49.1 and 30.6 nm, respectively (Figure 6A). Across a nine-setting sweep from *d* = 130 to 290 nm, ensemble peaks ranged from 360 to 756 nm and the standard deviation among realisations ranged from 16.7 to 54.5 nm (Figure 6B); the spread did not increase monotonically with spacing.

**Figure 6.**
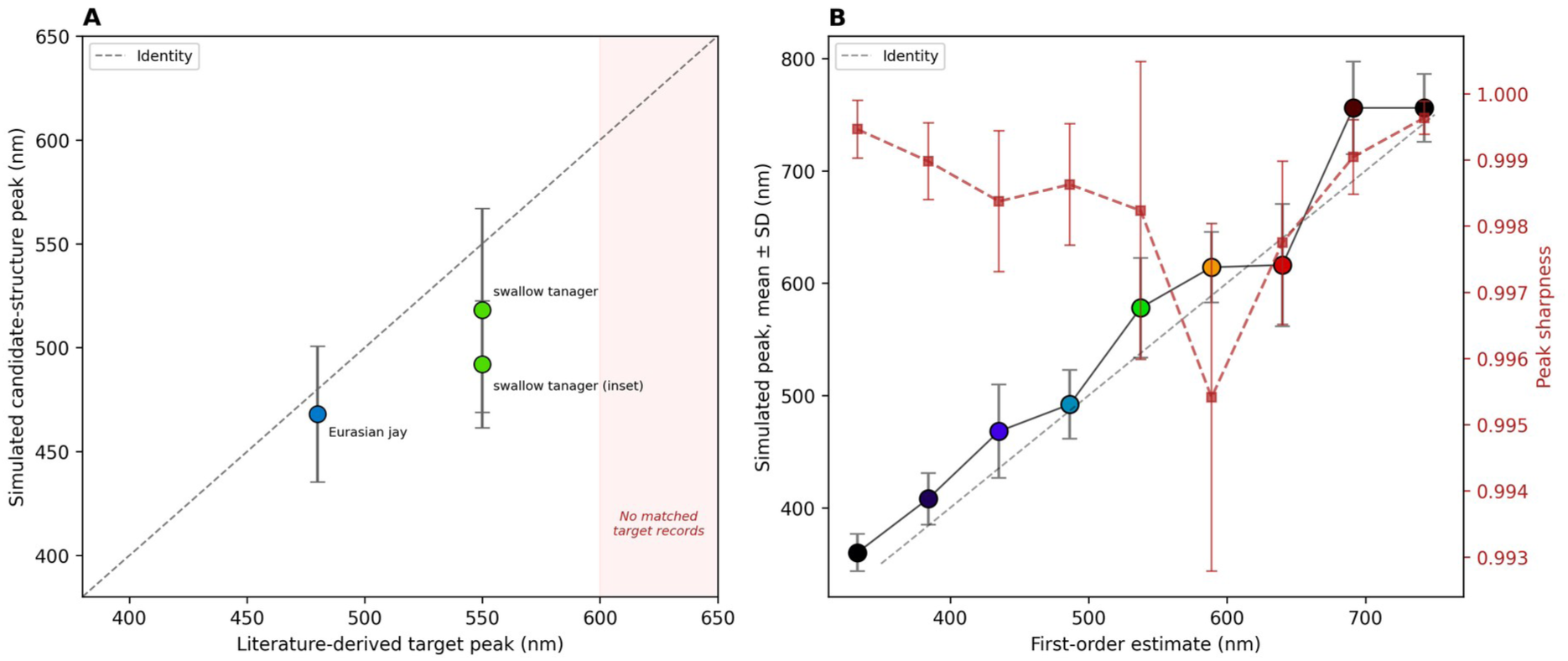
Variability among synthetic realisations. (A) Peak wavelength of the ensemble-averaged reflectance versus the literature-derived target peak for the three species/panel-linked candidate structures in Figure 5B. Error bars show the standard deviation of peak wavelengths across 20 independently generated structures with fixed model parameters; the shaded region denotes wavelengths without matched target records. (B) The same ensemble-spectrum peak and realisation-level standard deviation over a characteristic-spacing sweep from *d* = 130 to 290 nm. The red dashed series reports mean peak sharpness, with its standard deviation on the right-hand axis. For each realisation, peak sharpness is defined as the maximum of the peak-normalized spectrum minus its 25th percentile; it is a dimensionless contrast statistic, not a measure of spectral linewidth.

These repeated-realisation intervals isolate generator-driven variation because pore fraction, disorder and nominal spacing are fixed within each setting. A nominal candidate therefore corresponds to a distribution of simulated peak wavelengths rather than to one exact value, providing the uncertainty context for the 12–58 nm absolute differences in the record-linked comparisons.

A synthetic spacing series illustrated the first-order spacing–wavelength relation across 11 adjacent texture regions (Supplementary Figure S1). At a fixed effective refractive index of 1.28, the analytically predicted peak increased from 384 to 730 nm as the assigned characteristic spacing increased from 150 to 285 nm. These values were calculated from the first-order relation rather than extracted from the displayed textures. The next analysis examined the spacing-dependent trend in explicit three-dimensional volumes and a full-wave calculation. The biological mechanism linking quasi-ordered structure to reflected wavelength is summarized in Figure 7.

**Figure 7.**
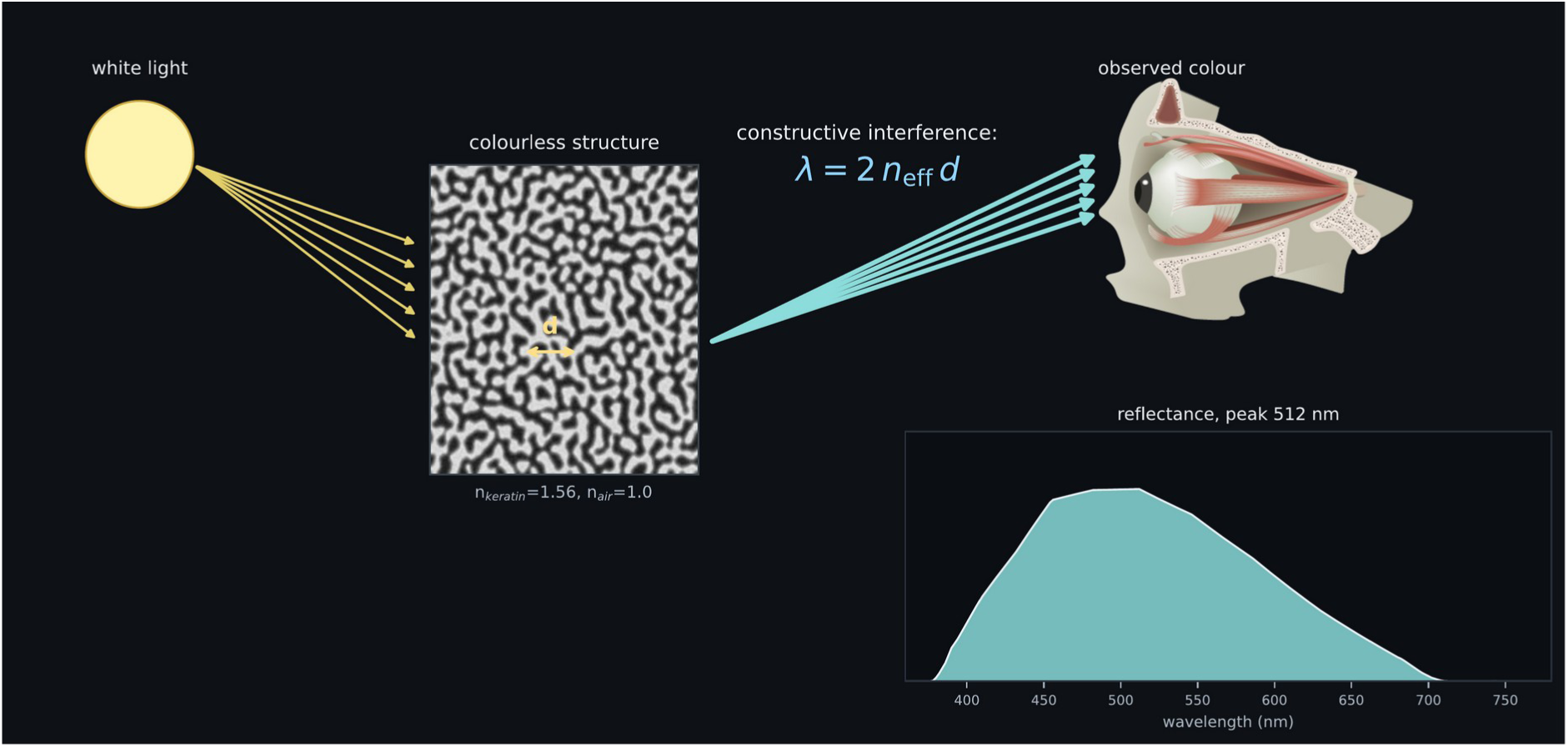
Physical mechanism underlying non-iridescent structural colour. Broadband illumination is selectively reflected by constructive interference within the quasi-ordered keratin-air nanostructure. The dominant wavelength depends on characteristic spacing *d* according to *λ ≈* 2 *n_eff_ d*. For the dominant back-scattering peak, *n_eff_* is the effective refractive index and *d* is the dominant structural correlation length (Noh et al. 2010, eq. 2). The eye image is from TogoTV (© 2016 DBCLS TogoTV, CC BY 4.0; https://creativecommons.org/licenses/by/4.0/deed.ja).

### 3.5 Three-dimensional and FDTD calculations

The next analysis examined whether the spacing-dependent response extended from two-dimensional sections to explicit three-dimensional geometry. Keratin–air voxel volumes with *d* = 160, 200 and 250 nm produced reflectance peaks at 460, 528 and 736 nm, respectively (Figure 8), compared with first-order estimates of 410, 512 and 640 nm; the absolute differences were 50, 16 and 96 nm.

**Figure 8.**
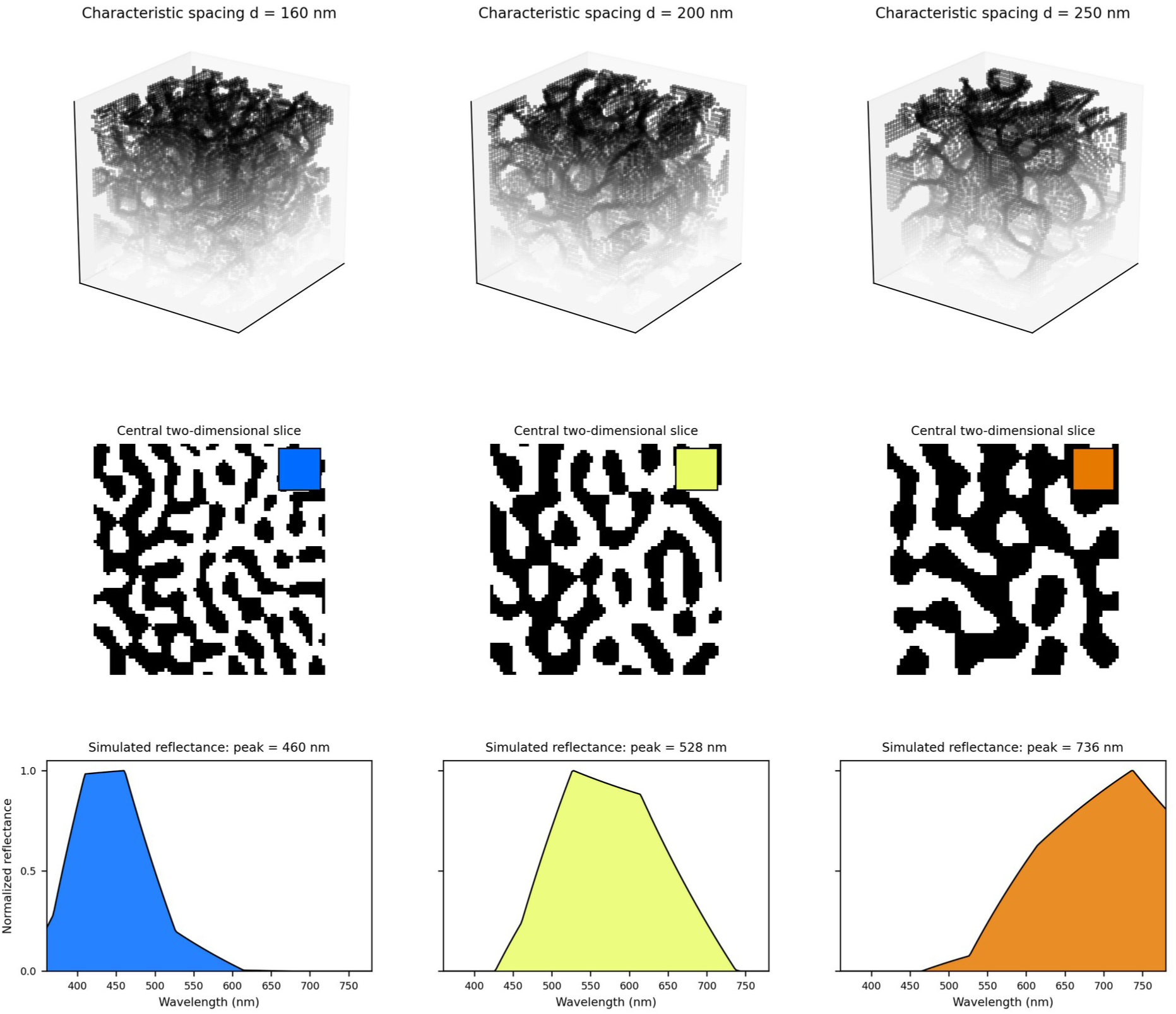
Three-dimensional structure-factor calculations. Columns show representative keratin-air voxel volumes, central slices, CIE-derived colour swatches and normalized spectra at *d* = 160, 200 and 250 nm. Calculations used pore fraction 0.50, disorder 0.12, 20 nm voxels and seed 1; spectra were computed from 72 × 72 × 72 voxel volumes and 40 × 40 × 40 voxel volumes are displayed. Peaks were 460, 528 and 736 nm; the first-order estimates were 410, 512 and 640 nm.

The three-dimensional calculation therefore reproduced the spacing-dependent red shift across all three structures. The deviation from the first-order estimate was smallest at *d* = 200 nm and increased to 96 nm for the *d* = 250 nm volume, showing where explicit geometry departed most strongly from the analytical estimate.

A full-wave MEEP FDTD analysis provided a preliminary cross-check of the spectral-region trend across resolution, smoothing and seed settings. For the *d* = 185 nm, seed-1 volume, peak wavelengths at resolutions of 24, 36 and 50 pixels/µm were 477, 381 and 576 nm without subpixel smoothing and 576, 581 and 473 nm with smoothing, spanning 381–581 nm across the six settings (Supplementary S5). At 24 pixels/µm with smoothing, three seeds gave peaks of 401, 360 and 362 nm for *d* = 140 nm (mean 374.3 nm; SD 19.1 nm) and 608, 670 and 663 nm for *d* = 250 nm (mean 647.0 nm; SD 27.6 nm). The two spacing groups remained separated, supporting the direction of the spacing-dependent shift, while the absolute peak at fixed *d* varied with resolution and smoothing. The FDTD result therefore remains a preliminary in-silico cross-check rather than a converged quantitative validation.

### 3.6 The idealized multilayer model produced an angle-dependent blue shift

The angle response of an ordered iridescent mechanism was examined with the idealized multilayer calculation. Increasing the incidence angle progressively shifted the reflectance spectrum towards shorter wavelengths (Figure 9B): the peak moved from 478 nm at 0° to 472, 456, 434 and 408 nm at 15°, 30°, 45° and 60°, respectively, for a total blue shift of 70 nm (Figure 9C). The CIE-derived display colour changed with the same trend, from cyan at normal and low incidence angles to blue at 45° and violet at 60°. Thus, the ordered multilayer produced the explicit angle-dependent blue shift expected for an iridescent structure.

**Figure 9.**
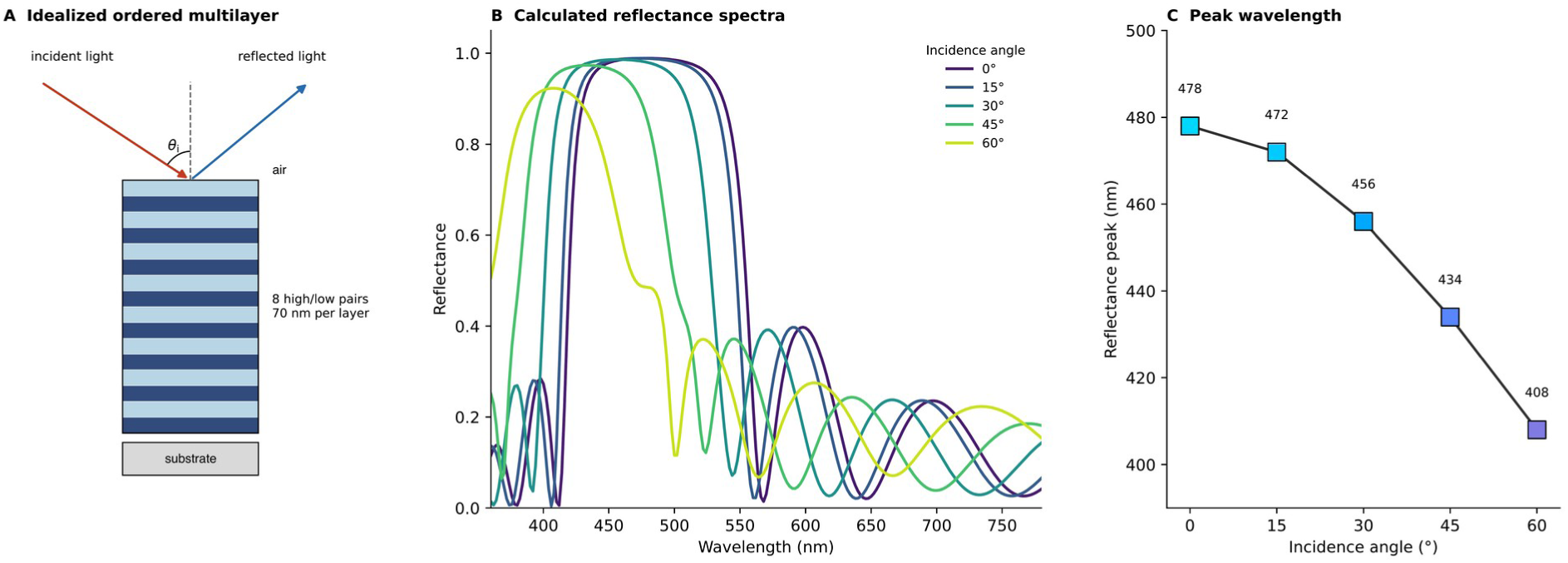
Angle-dependent reflectance of an idealized ordered multilayer. (A) Eight high/low-index pairs (n = 2.0/1.40; 70 nm layers) between air (n = 1.0) and a substrate (n = 1.5); θi is the incidence angle. (B) Unpolarized transfer-matrix spectra at 0°, 15°, 30°, 45° and 60°. (C) Peak wavelength and CIE-derived sRGB at each angle. The 478-to-408 nm shift illustrates the expected blue shift for an ordered iridescent reference mechanism.

### 3.7 Simulated spectra produced corresponding colour visualizations

Calculated spectra were then converted to CIE-derived sRGB to provide an interpretable visual output. Across the synthetic spacing series, the reflectance peak shifted from 390 nm at *d* = 140 nm to 744 nm at *d* = 275 nm, and the corresponding colour patches progressed from blue and cyan through green to yellow and orange (Figure 10A,B). Each tinted structure and uniform patch was generated from the same simulated spectrum, directly linking the spacing-dependent optical response to its calculated display colour.

**Figure 10.**
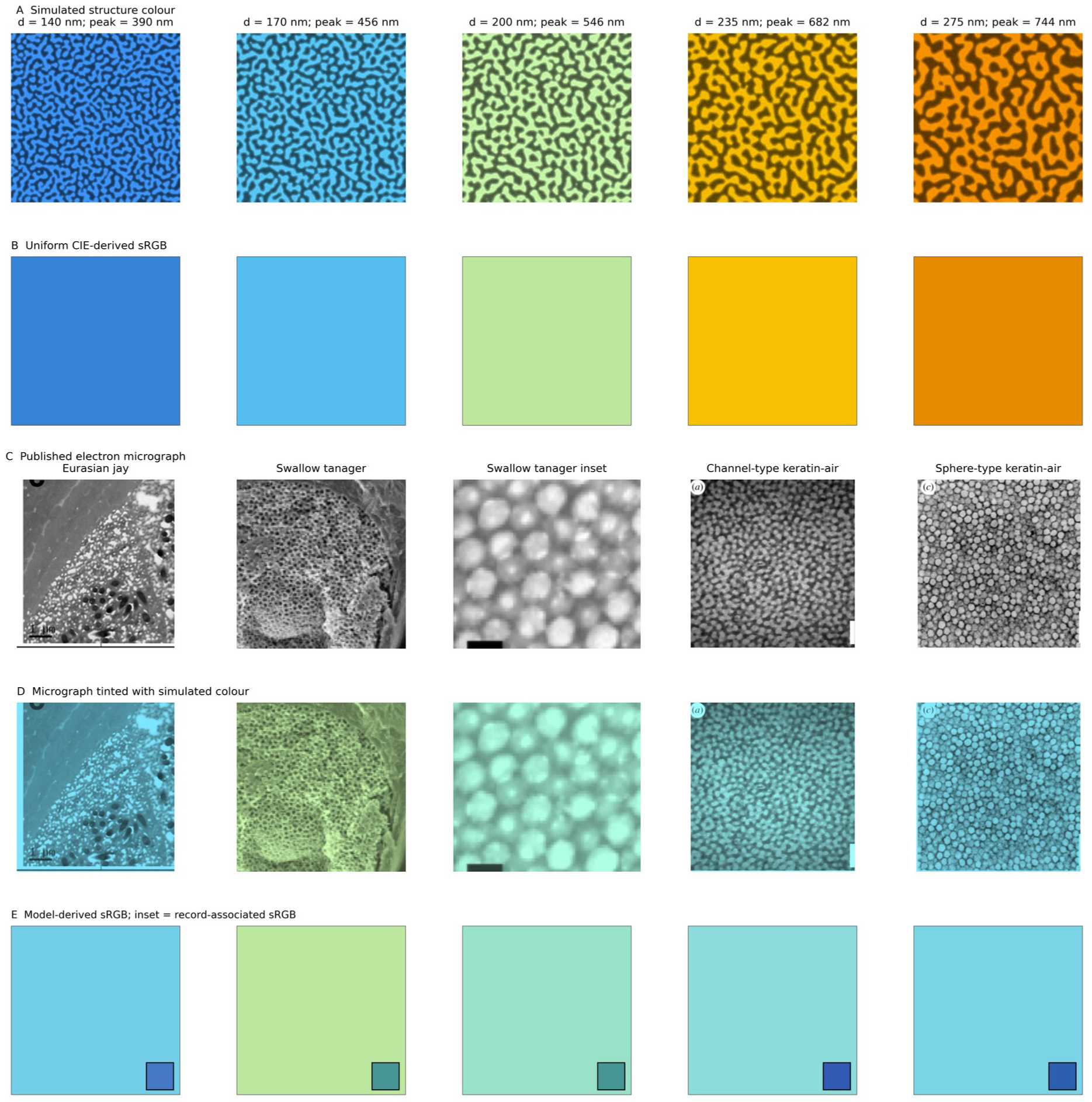
Colour visualizations derived from simulated reflectance spectra. (A) Synthetic structures with characteristic spacings *d* = 140, 170, 200, 235 and 275 nm, tinted using the CIE-derived sRGB value of their simulated spectra; the corresponding peaks shift from 390 to 744 nm. (B) Uniform sRGB patches generated directly from the same spectra, showing a progression from blue and cyan through green to yellow and orange. (C) Published electron micrographs used as literature-linked keratin-air examples. (D) The micrographs tinted with the simulated colour obtained from each declared candidate spacing while preserving grayscale image intensity. (E) Uniform model-derived sRGB patches; inset squares show the separate record-associated sRGB annotations. Panels C-E provide a qualitative visual comparison, while quantitative wavelength analyses are presented in Figures 4-6. Published micrographs are reproduced from Parnell et al. (2015; CC BY 4.0), Bazzano et al. (2020; bioRxiv preprint, CC BY), and Prum et al. (2009; CC BY 2.0).

The same colour mapping was finally applied to five literature-linked keratin–air candidates using their declared spacings, allowing model-derived colours to be compared with record-associated visual annotations (Figure 10C–E). The model-derived colours were predominantly cyan to green, whereas several record-associated sRGB insets were bluer. This comparison identifies the direction of the visual difference; the quantitative peak-wavelength results that explain its magnitude are given in Figures 4–6.

## 4. Discussion

The results support a bounded route from avian nanostructure to computational structural-colour design. Literature-informed spacings were used to generate synthetic structures, and coherent single-scattering theory predicted their spectra without fitting the displayed target colours. Given the heterogeneous target provenance and seed-dependent variation (§3.4, Figure 6), the agreement is interpreted as in-silico consistency between structure and colour rather than independent experimental validation.

These findings are consistent with previous studies showing that quasi-ordered photonic nanostructures behave as amorphous photonic materials (Noh et al. 2010; Saranathan et al. 2012). The present framework extends that physical relationship into a prior-constrained workflow that links biological nanostructures to predicted colour and desired colours to candidate nanostructures. Kingfisher feathers provide a well-characterized biological example in which pigments, spongy nanostructures and thin-film effects jointly contribute to colour (Stavenga et al. 2011b).

### From descriptive biology to structural-colour design

The principal contribution of Kawasemi Colours is not a new optical theory, but an open, provenance-tracked implementation that connects published avian nanostructures to interpretable forward prediction, prior-constrained inverse design and physics-based in-silico checks within a single reproducible workflow. Applied to the curated records, the workflow produced lower prediction errors for mechanism-matched keratin–air structures than for out-of-domain melanosome arrays, while controlled two- and three-dimensional simulations reproduced the expected spacing-dependent spectral shift. These results support the internal consistency and defined applicability of the framework, but do not constitute independent experimental validation. Its inverse component does not search all possible optical structures; it explores candidate geometries within the keratin–air pore architectures found in feathers and evaluates how spacing, pore fraction and order alter the simulated optical response. This defined design space preserves the biological question at the centre of the study: how variation within a feather-like pore network can generate colour variation.

Colour-only inverse problems are intrinsically non-unique because different structures and materials can yield similar colours. I therefore report one prior-constrained keratin–air candidate together with a model-conditional spacing range, rather than a unique solution. This restricted formulation is consistent with the general inverse-design literature (Khaireh-Walieh et al. 2023) while keeping the inference within the biological and material assumptions tested here.

Electron micrographs of structurally coloured feathers remain scattered across the literature and are rarely accompanied by reusable metadata or computational tools. Kawasemi Colours therefore releases curated metadata, reproducible pipelines and interactive applications with source-level provenance for image scale, imaging conditions and colour or peak-wavelength measurements. Future records can broaden the represented avian architectures, test generalizability and refine predictive models when supported by appropriate validation.

### Limitations

The real-image dataset comprises 26 avian micrographs and five biomimetic comparison images. The avian records include 14 named species and four panels without species-level identification in the current metadata. The measured-scale subset contains six records and has larger exploratory errors than the heuristic-scale subset (Supplementary Table S3), although morphological differences between the subsets confound the effect of calibration on prediction error. Target peaks for the real-image records span 418–599 nm and are variously literature-reported, label-derived or author-derived (Supplementary Table S2).

The analytical model is based on coherent single scattering and therefore neglects multiple scattering, near-field interactions and wavelength-dependent material dispersion. These effects are expected to become increasingly important for highly disordered or strongly scattering structures.

Colour prediction from a single two-dimensional electron micrograph is inherently approximate because any section represents only a partial view of the underlying three-dimensional nanostructure. The accompanying three-dimensional analyses preserve the overall spacing-dependent trend, but precise quantitative prediction ultimately requires volumetric structural information.

The present FDTD calculations use a periodically repeated lateral cell spanning approximately 2.9 structural periods, and peak estimates vary across resolution, smoothing and seed settings (Supplementary S5). Their evidential scope is qualitative agreement in spectral region and spacing-dependent trend. Quantitative full-wave benchmarking will require larger finite-domain volumes, broader disorder-realisation sampling and systematic convergence analysis.

### Outlook

Future work should prioritize three-dimensional reconstruction from electron tomography, quantitative FDTD benchmarking, larger openly licensed datasets, prediction of angle- and polarization-dependent reflectance and experimental validation. The inverse-design outputs define candidate structural descriptors, but translating them into controllable fabrication variables will require material-specific calibration and validation with fabricated photonic materials. These steps will determine whether the present structure–colour relationships can support predictive engineering beyond the current in-silico framework.

## 5. Acknowledgements and declarations

### Author contributions

Masaki Suimye Morioka: conceptualization, software implementation, formal analysis, visualization, and original draft preparation. Anthropic Claude Code and OpenAI Codex were used, under the author’s direction and verification, for implementation assistance and manuscript revision. The author is responsible for all analyses, interpretations, and the final manuscript.

### Data and code availability

Source code, interactive web applications, analysis and validation pipelines, and figure-generation scripts are available under the MIT License at https://github.com/suimye/kawasemi-colours. The versioned release is archived on Zenodo (Morioka 2026; doi:10.5281/zenodo.23445928).

Author-created metadata and synthetic structures are provided under CC BY 4.0. Third-party images retain their original licences and attribution requirements; the software licence does not apply to them. Image-level provenance and redistribution status are supplied in Supplementary Tables S1–S2. The versioned code and data archive is identified above; the corrected supplementary licensing table supersedes blanket licence descriptions for third-party images.

Restricted or unconfirmed image files are not reproduced in this manuscript. Their source publications, figure identifiers and redistribution status are documented in Supplementary Tables S1–S2. Reproduction of analyses that use those images requires obtaining the original inputs subject to applicable permissions. The public GitHub repository and versioned Zenodo archive provide the analysis software.

