## supplemental file for "Kawasemi-Colours: an open end-to-end platform for structure–colour design based on avian structural colour"

Masaki Suimye Morioka

Database Division for Life Science (DBCLS)
BioData Science Initiative (BSI), National Institute of Genetics
Research Organization of Information and Systems

Code: https://github.com/suimye/kawasemi-colours

Versioned archive (v0.1.1): https://doi.org/10.5281/zenodo.23445928

#### Contents

Supplementary Figure S1. Synthetic spacing gradient
Supplementary Tables S1–S2. Licensing and record provenance
Supplementary Table S3. Scale-stratified validation
Supplementary Table S4. Validation populations and protocol
Supplementary S5. FDTD reproducibility and limitations

### Supplementary Figure S1

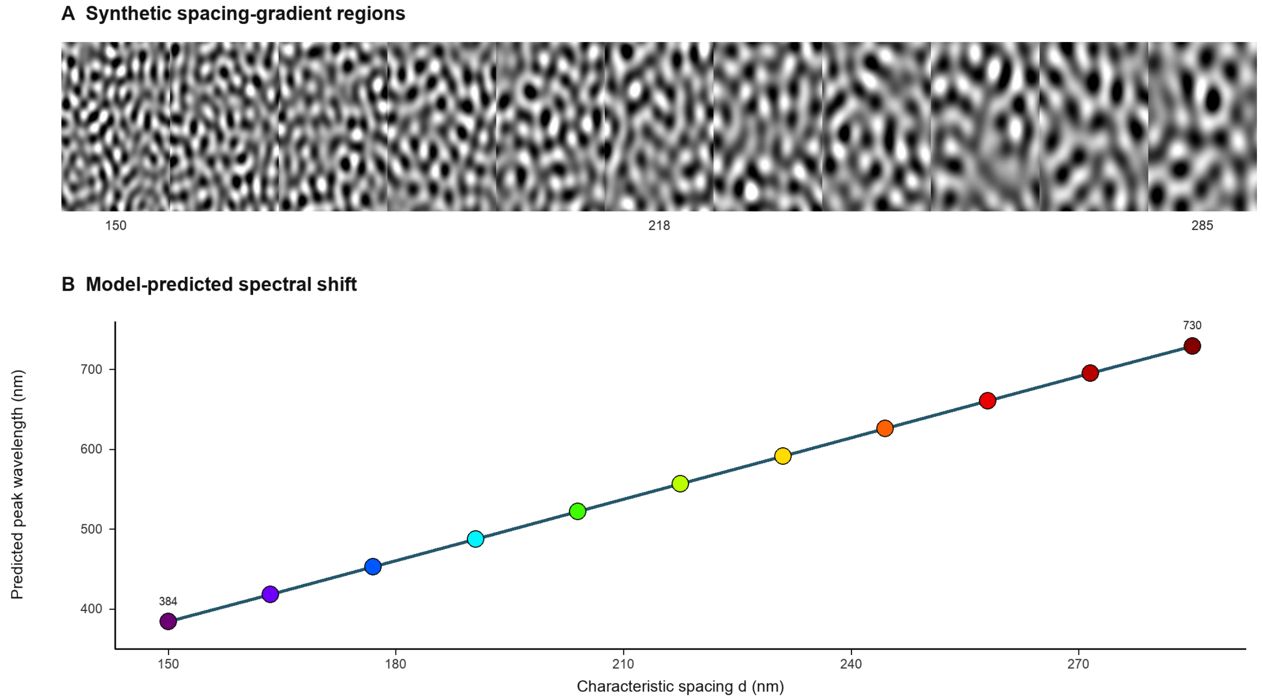


Analytically predicted peak shift across a synthetic spacing series. Eleven independently generated texture regions are placed side by side, with assigned characteristic spacing d increasing from 150 to 285 nm while pore fraction (0.50), disorder treatment and refractive indices are held fixed. The plotted wavelengths are calculated from the first-order coherent-scattering relation $\lambda_0 = 2\, n_{\mathrm{eff}}\, d$ with $n_{\mathrm{eff}} = 1.28$, shifting from 384 to 730 nm; they are not reflectance peaks extracted from the displayed textures. Point colours are display aids derived from wavelength and are not measured specimen colours. This illustration shows the direction of the first-order spacing-dependent shift, not an independently validated continuous-structure simulation.

#### Supplementary Table S1. Image-level licensing and provenance

This table inventories the 31 curated real-image records. Analysis eligibility is specified separately in Supplementary Tables S3–S4. An article-level open licence does not automatically cover third-party figures. The original image credit line controls the redistribution assessment.

Nine review-derived records have permission credits rather than an established image-level CC BY licence. They are excluded from the redistributable image subset unless separate permission is documented. Their analysis records and the reported numerical results are retained. Six of these images have been removed from Figure 2. This table makes no blanket legal assertion that non-commercial status alone permits machine-learning use.

CC BY images retain the version and attribution requirements of their original source. CC BY-NC images are not unrestricted redistributable material. Restricted or unconfirmed images must be obtained from the original publication subject to applicable permissions. Historical public archives require a separate file-level review; this corrected table does not establish that an earlier archive has been updated.

Full source URLs, analysis-use fields and image-specific notes are supplied in Supplementary_Table_S1_data_licenses.csv. For the biomimetic particle panels, the original source is Kohri et al. (2019), doi:10.1021/acs.langmuir.9b00400. For the Steller's-jay micrograph, it is Shawkey and Hill (2006), doi:10.1242/jeb.02115.

| Record ID | Licence / credit | Redistribution |
|---|---|---|
| PMC9832292_fig2b_blue | CC BY 4.0 | Yes (with attribution) |
| PMC9832292_fig2b_green | CC BY 4.0 | Yes (with attribution) |
| PMC9832292_fig3a_blue_etch60 | CC BY 4.0 | Yes (with attribution) |
| PMC9832292_fig3a_blue_etch120 | CC BY 4.0 | Yes (with attribution) |
| PMC8599086_fig1c_bluevane | CC BY-NC 4.0 | Non-commercial only, with attribution; excluded from unrestricted subset |
| PMC8599086_fig1d_whitedot | CC BY-NC 4.0 | Non-commercial only, with attribution; excluded from unrestricted subset |
| PMC8505553_fig4_rockdove | CC BY (original source credit) | Yes (with attribution) |
| PMC8505553_fig4_stellersjay | Third-party credit; review CC BY does not establish image rights | Excluded; separate permission not documented |
| PMC8505553_fig5a_parotia | Third-party credit; review CC BY does not establish image rights | Excluded; separate permission not documented |
| PMC8505553_fig5b_teal | CC BY (original source credit) | Yes (with attribution) |
| PMC8505553_fig5c_starling | Third-party credit; review CC BY does not establish image rights | Excluded; separate permission not documented |
| PMC8505553_fig5d_racket | Third-party credit; review CC BY does not establish image rights | Excluded; separate permission not documented |
| PMC8505553_fig6d_sphere | Third-party credit; review CC BY does not establish image rights | Excluded; separate permission not documented |
| PMC8505553_fig6d_roundish | Third-party credit; review CC BY does not establish image rights | Excluded; separate permission not documented |
| PMC8505553_fig6d_mixed | Third-party credit; review CC BY does not establish image rights | Excluded; separate permission not documented |
| PMC8505553_fig6d_ellipsoid | Third-party credit; review CC BY does not establish image rights | Excluded; separate permission not documented |
| PMC8505553_fig6d_elong | Third-party credit; review CC BY does not establish image rights | Excluded; separate permission not documented |

| Record ID | Licence / credit | Redistribution |
|---|---|---|
| STAVENGA2011_king_fig3F | not CC (green-OA) | No |
| BIORXIV_tersina_fig6c | CC BY | Yes (with attribution) |
| BIORXIV_tersina_fig6b_inset | CC BY | Yes (with attribution) |
| BIORXIV_tersina_fig6d_inset | CC BY | Yes (with attribution) |
| PMC2706469_fig1e_channel | CC BY 2.0 | Yes (with attribution) |
| PMC2706469_fig1a_channel | CC BY 2.0 | Yes (with attribution) |
| PMC2706469_fig1c_sphere | CC BY 2.0 | Yes (with attribution) |
| PMC4685390_fig6c_jay | CC BY 4.0 | Yes (with attribution) |
| PMC4685390_fig6d_jay | CC BY 4.0 | Yes (with attribution) |
| PMC4685390_fig6e_jay | CC BY 4.0 | Yes (with attribution) |
| PMC3427513_fig2d_broadbill | not CC (c) Royal Society | No |
| PMC3427513_fig2e_bluebird | not CC (c) Royal Society | No |
| PMC3427513_fig2f_cotinga | not CC (c) Royal Society | No |
| PMC2706469_fig2f_sphere | CC BY 2.0 | Yes (with attribution) |

#### Supplementary Table S2. Record-level provenance summary

Identity and measurement fields are presented in two linked tables, using the same record IDs. Complete provenance fields are supplied in Supplementary_Table_S2_provenance.csv. Verify denotes a source value requiring manual checking before it is treated as a direct spectral measurement; Author-derived denotes a value derived from a source description; Display only is not a spectral measurement.

| Record ID | Type | Species / material | Structure | Source panel |
|---|---|---|---|---|
| PMC9832292_fig2b_blue | image | Phasianus colchicus | melanosome array | Figure 2b (blue barbule) |
| PMC9832292_fig2b_green | image | Phasianus colchicus | melanosome array | Figure 2b (green barbule) |
| PMC9832292_fig3a_blue_etch60 | image | Phasianus colchicus | melanosome array | Figure 3a-ii (blue barbule, 60 s etch) |
| PMC9832292_fig3a_blue_etch120 | image | Phasianus colchicus | melanosome array | Figure 3a-iii (blue barbule, 120 s etch) |
| PMC9832292_fig3d_green_etch60 | proxy | Phasianus colchicus | melanosome array | Figure 3d-ii (green barbule, 60 s etch) |
| PMC9832292_fig3d_green_etch120 | proxy | Phasianus colchicus | melanosome array | Figure 3d-iii (green barbule, 120 s etch) |
| PMC11296197_fig1b_crown | proxy | Cyanistes caeruleus | spongy | Figure 1b (TEM spongy nanostructure) |
| PMC8599086_fig1c_bluevane | image | Cyanistes caeruleus | spongy | Figure 1C (SEM blue vane cross-section) |
| PMC8599086_fig1d_whitedot | image | Cyanistes caeruleus | spongy | Figure 1D (SEM white dot cross-section) |
| PMC10032483_fig2c_barbule | proxy | Pica serica | melanosome array | Figure 2C (TEM hexagonal melanin stack) |
| PMC10032483_fig2c_violet_domain | proxy | Pica serica | melanosome array | Figure 2C (TEM, violet domain) |
| PMC10032483_fig2c_blue_domain | proxy | Pica serica | melanosome array | Figure 2C (TEM, blue domain) |
| PMC8505553_fig4_rockdove | image | Columba livia | melanosome array | Figure 4 (rock dove barbule) |
| PMC8505553_fig4_stellersjay | image | Cyanocitta stelleri | spongy | Figure 4 (Steller's jay blue barb) |
| PMC8505553_fig5a_parotia | image | Parotia lawesii | melanosome array | Figure 5a (Lawes' parotia breast) |
| PMC8505553_fig5b_teal | image | Anas crecca | melanosome array | Figure 5b (green-winged teal wing) |
| PMC8505553_fig5c_starling | image | Cinnyricinclus leucogaster | melanosome array | Figure 5c (violet-backed starling) |
| PMC8505553_fig5d_racket | image | Ocreatus underwoodii | melanosome array | Figure 5d (racket-tail hummingbird gorget) |
| PMC10962638_fig2_spongy | proxy | Malurus leucopterus | spongy | Figure 2 (barb cross-section schematic: cortex/spongy/melanosome) |
| PMC8505553_cotinga_spongy | proxy | Cotinga maynana | spongy | mechanism ref (coherent scattering, Fig 1e) |
| PMC8505553_bluejay_spongy | proxy | Cyanocitta cristata | spongy | mechanism ref (amorphous nanostructure -> non-iridescent) |
| PMC8505553_bluebird_spongy | proxy | Sialia sialis | spongy | mechanism ref (coherent scattering) |

| Record ID | Type | Species / material | Structure | Source panel |
|---|---|---|---|---|
| SPECIESREF_alcedo_spongy | proxy | Alcedo atthis | spongy | species reference (no OA SEM figure) |
| PMC8505553_fig6d_sphere | image | biomimetic (synthetic melanin nanoparticles) | quasi ordered sphere | Figure 6d (sphere) |
| PMC8505553_fig6d_roundish | image | biomimetic (synthetic melanin nanoparticles) | quasi ordered sphere | Figure 6d (roundish) |
| PMC8505553_fig6d_mixed | image | biomimetic (synthetic melanin nanoparticles) | quasi ordered sphere | Figure 6d (mixed) |
| PMC8505553_fig6d_ellipsoid | image | biomimetic (synthetic melanin nanoparticles) | quasi ordered sphere | Figure 6d (ellipsoid) |
| PMC8505553_fig6d_elong | image | biomimetic (synthetic melanin nanoparticles) | quasi ordered sphere | Figure 6d (elong) |
| STAVENGA2011_king_fig3F | image | Alcedo atthis | spongy | Fig 3F (tail barb spongy, 2um) |
| BIORXIV_tersina_fig6c | image | Tersina viridis | spongy | Fig 6c (SEM spongy matrix) |
| BIORXIV_tersina_fig6b_inset | image | Tersina viridis | quasi ordered sphere | Fig 6b inset (high-mag sphere-type air array) |
| BIORXIV_tersina_fig6d_inset | image | Tersina viridis | quasi ordered sphere | Fig 6d inset (high-mag sphere-type array) |
| PMC2706469_fig1e_channel | image | Aves (Prum 2009, channel-type) | channel | Fig 2e (high-mag channel, 120nm bar, hi-res) |
| PMC2706469_fig1a_channel | image | Aves (Prum 2009, channel-type) | channel | Fig 2a (channel beta-keratin, hi-res) |
| PMC2706469_fig1c_sphere | image | Aves (Prum 2009, sphere-type) | quasi ordered sphere | Fig 2c (sphere beta-keratin, hi-res) |
| PMC4685390_fig6c_jay | image | Garrulus glandarius | channel | Fig 6c (channel nanostructure, 1um) |
| PMC4685390_fig6d_jay | image | Garrulus glandarius | channel | Fig 6d (channel nanostructure, 1um) |
| PMC4685390_fig6e_jay | image | Garrulus glandarius | channel | Fig 6e (TEM channel nanostructure, 1um) |
| PMC3427513_fig2d_broadbill | image | Serilophus lunatus | channel | Fig 2d (SEM channel nanostructure) |
| PMC3427513_fig2e_bluebird | image | Sialia sialis | channel | Fig 2e (SEM channel nanostructure) |
| PMC3427513_fig2f_cotinga | image | Cotinga maynana | quasi ordered sphere | Fig 2f (SEM/TEM sphere nanostructure) |
| PMC2706469_fig2f_sphere | image | Aves (Prum 2009, sphere-type) | quasi ordered sphere | Fig 2f (ordered sphere array, hi-res) |

#### Supplementary Table S2 continued — values and calibration

| Record ID | Peak (nm) | Period (nm) | Scale (px/µm) | Value status |
|---|---|---|---|---|
| PMC9832292_fig2b_blue | 480 | — | 106.7 | Verify |
| PMC9832292_fig2b_green | 520 | — | 110.0 | Verify |
| PMC9832292_fig3a_blue_etch60 | 430 | — | 315.0 | Verify |
| PMC9832292_fig3a_blue_etch120 | 418 | — | 315.0 | Verify |
| PMC9832292_fig3d_green_etch60 | 497 | — | Uncalibrated | Author-derived |
| PMC9832292_fig3d_green_etch120 | 485 | — | Uncalibrated | Author-derived |
| PMC11296197_fig1b_crown | 440 | — | Uncalibrated | Author-derived |
| PMC8599086_fig1c_bluevane | 470 | — | Uncalibrated | Verify |
| PMC8599086_fig1d_whitedot | — | — | Uncalibrated | No peak |
| PMC10032483_fig2c_barbule | 520 | — | Uncalibrated | Author-derived |
| PMC10032483_fig2c_violet_domain | 420 | — | Uncalibrated | Author-derived |
| PMC10032483_fig2c_blue_domain | 460 | — | Uncalibrated | Author-derived |
| PMC8505553_fig4_rockdove | 530 | — | Uncalibrated | Verify |
| PMC8505553_fig4_stellersjay | 460 | — | Uncalibrated | Verify |
| PMC8505553_fig5a_parotia | 470 | — | Uncalibrated | Verify |
| PMC8505553_fig5b_teal | 530 | — | 84.0 | Verify |
| PMC8505553_fig5c_starling | 420 | — | Uncalibrated | Verify |
| PMC8505553_fig5d_racket | 540 | — | 138.0 | Verify |
| PMC10962638_fig2_spongy | 470 | — | Uncalibrated | Author-derived |
| PMC8505553_cotinga_spongy | 490 | — | Uncalibrated | Author-derived |
| PMC8505553_bluejay_spongy | 470 | — | Uncalibrated | Author-derived |
| PMC8505553_bluebird_spongy | 475 | — | Uncalibrated | Author-derived |
| SPECIESREF_alcedo_spongy | 485 | — | Uncalibrated | Author-derived |
| PMC8505553_fig6d_sphere | 599.4 | — | Uncalibrated | Verify |
| PMC8505553_fig6d_roundish | 580.8 | — | Uncalibrated | Verify |
| PMC8505553_fig6d_mixed | 573.4 | — | Uncalibrated | Verify |
| PMC8505553_fig6d_ellipsoid | 487.8 | — | Uncalibrated | Verify |
| PMC8505553_fig6d_elong | 480.4 | — | Uncalibrated | Verify |
| STAVENGA2011_king_fig3F | 485 | — | Uncalibrated | Verify |
| BIORXIV_tersina_fig6c | 550 | — | Uncalibrated | Display only |
| BIORXIV_tersina_fig6b_inset | 550 | — | Uncalibrated | Display only |
| BIORXIV_tersina_fig6d_inset | 550 | — | Uncalibrated | Display only |
| PMC2706469_fig1e_channel | 470 | — | Uncalibrated | Verify |
| PMC2706469_fig1a_channel | 470 | — | Uncalibrated | Verify |
| PMC2706469_fig1c_sphere | 460 | — | Uncalibrated | Verify |
| PMC4685390_fig6c_jay | 480 | 150 | Uncalibrated | Verify |
| PMC4685390_fig6d_jay | 490 | 180 | Uncalibrated | Verify |

| Record ID | Peak (nm) | Period (nm) | Scale (px/µm) | Value status |
|---|---|---|---|---|
| PMC4685390_fig6e_jay | 480 | 175 | Uncalibrated | Verify |
| PMC3427513_fig2d_broadbill | 470 | — | Uncalibrated | Verify |
| PMC3427513_fig2e_bluebird | 465 | — | Uncalibrated | Verify |
| PMC3427513_fig2f_cotinga | 485 | — | Uncalibrated | Verify |
| PMC2706469_fig2f_sphere | 460 | — | Uncalibrated | Verify |

#### Supplementary Table S3. Scale-stratified validation of FEATS_V1 prediction

Length-valued FEATS_V1 descriptors were calculated with the record-specific scale.px_per_um only when the record was marked calibrated: true. All other records retain the extractor's explicit heuristic conversion and are reported as the **heuristic-scale** subset. The latter is not an absolute nanometre measurement. Models were trained as described in model/validate_real_grouped.py: 800 synthetic structures plus the available non-held-out records. Evaluation is restricted to 28 real_image=true records eligible for colour-model training. The three swallow-tanager records are excluded because their retained sRGB values are illustrative display swatches, not spectrophotometrically calibrated colour targets. Results are exploratory because the measured-scale subset contains only six records and still spans different optical morphologies.

| Validation split | All eligible real-image records (n=28) | Measured-scale records (n=6) | Heuristic-scale records (n=22) |
|---|---|---|---|
| Random 5-fold MAE, median (nm) | 58.9, 50.1 | 90.1, 85.3 | 50.4, 37.7 |
| Leave-one-species-out MAE, median (nm) | 55.7, 40.1 | 87.9, 73.4 | 47.0, 33.5 |
| Leave-one-paper-out MAE, median (nm) | 57.0, 41.1 | 87.9, 72.5 | 48.6, 36.7 |

The poorer measured-scale-subset values do not establish that calibrated scales reduce performance. They show instead that the present data are too small and too heterogeneous to make a reliable absolute-scale performance claim. In particular, the six calibrated records include several melanosome-array examples outside the quasi-ordered keratin–air mechanism targeted by the primary simulator. Future evaluation should use an independently acquired, scale-calibrated, non-iridescent keratin–air test set with reflectance spectra measured under recorded geometry.

#### Supplementary Table S4. Validation populations, randomization, and synthetic training protocol

All results below use the eligible record set after excluding the three swallow-tanager records with non-calibrated illustrative sRGB display swatches. The code and fixed seeds are stated so that the distinct evaluation populations are not conflated.

| Analysis | Evaluated records | Training data in each fold | Split and seed | Current result |
|---|---|---|---|---|
| train_real.py random 5-fold | 39 eligible records: 28 real_image=true and 11 synthetic-proxy metadata records; mixed optical morphologies | 800 on-the-fly synthetic structures plus the non-held-out eligible records | np.random.default_rng(0), five contiguous array_split folds after permutation | Peak MAE 57.2 nm; median 37.0 nm; saturation MAE 0.189 |
| validate_real_grouped.py random 5-fold | 28 eligible real_image=true records; mixed optical morphologies | 800 on-the-fly synthetic structures, all 11 proxies, and non-held-out real images | np.random.default_rng(0), five contiguous array_split folds after permutation | Peak MAE 58.9 nm; median 50.1 nm |
| validate_real_grouped.py LOSO | Same 28 real images, 16 species groups | 800 on-the-fly synthetic structures, all proxies, and records outside the held-out species | one held-out species at a time; no stochastic split beyond synthetic seed 0 | Peak MAE 55.7 nm; median 40.1 nm |
| validate_real_grouped.py LOPO | Same 28 real images, 7 source-paper groups | 800 on-the-fly synthetic structures, all proxies, and records outside the held-out source paper | one held-out source paper at a time; no stochastic split beyond synthetic seed 0 | Peak MAE 57.0 nm; median 41.1 nm |

The exact real-image random-five-fold assignment is archived in Supplementary_Table_S4_validation_folds_seed0.csv. These are exploratory pooled results: the evaluated set includes quasi-ordered keratin–air structures, melanosome arrays, and biomimetic controls. It is not a single-mechanism accuracy estimate.

#### On-the-fly synthetic training structures

The 11 synthetic-proxy metadata records above are not the synthetic training set. Each run generates 800 synthetic 96 × 96-pixel images in memory via model/synth.py, with generator seed 0. For each image, pore fraction is sampled uniformly from 0.30–0.70, disorder from 0.05–0.75, anisotropy from 0–0.6, and a target peak is sampled across the visible band before converting it to a characteristic spacing clipped to 90–320 nm. Pixel scale is independently sampled at 0.75–1.35 times DEFAULT_PX_PER_UM.

#### Interpretation of Figure 6

The 20-run dispersion is **synthetic-realisation variability**: only the random seed changes while the generator, spacing, pore fraction, disorder, image size, and optical model remain fixed. It does not include uncertainty from imaging, biological structural heterogeneity, target-label provenance, reflectance measurement, or model misspecification. It is therefore neither an experimental uncertainty interval nor a prediction interval for a biological specimen.

#### Supplementary S5 — FDTD reproducibility, convergence and boundary-condition caveats

This supplement documents the 3-D finite-difference time-domain (FDTD) runs referenced in Methods §2.8 and Results §3.5, including their software environment, exact commands, geometry and boundary conditions, resolution and smoothing sensitivity, and seed-specific results. The runs are retained only as a preliminary full-Maxwell cross-check: they are neither independent nor quantitative validation because the reported peaks are not numerically converged and the small structure is laterally periodic.

##### S5.1 Software and environment

| Component | Version |
|---|---|
| MEEP (pymeep, conda-forge) | 1.34.0 |
| Python | 3.11.15 |
| NumPy | 2.4.6 |
| SciPy | 1.17.1 |
| OS / arch | macOS (Darwin 24.6.0), arm64 (Apple Silicon) |

Installed via:

conda create -n meep -c conda-forge pymeep python=3.11

conda activate meep

##### S5.2 Structure generation (fixed across all runs unless stated)

The 3-D keratin/air voxel volume is generated by model/structure3d.py:

structure3d.generate_volume(d_nm, pore_fraction=0.5, disorder=0.12,

nm_per_vox=22.0, size=vox, seed=seed)

d_nm is the target characteristic spacing, size=vox is the number of voxels per side (24 throughout, i.e. a 24×24×24 volume, 22 nm/voxel → 528 nm per side, **≈2.9 periods per side** for d = 185 nm). seed selects one specific quasi-random disorder realisation; the committed script (model/fdtd3d_meep.py) exposes this as --seed (default 1), and exposes the MaterialGrid sensitivity setting as --subpixel-smoothing; the multi-seed and smoothing runs in §S5.5 can therefore be reproduced directly.

##### S5.3 MEEP simulation setup (model/fdtd3d_meep.py)

- **Cell**: sx = sy = L, sz = L + 2*(dpml + pad), where L = structure side in MEEP units (a = 1000 nm), dpml = 1.0 (a-units, i.e. 1000 nm), pad = 0.6 (600 nm). For the standard d = 185 nm / vox = 24 / 22 nm-per-vox case, L = 0.528 a (528 nm), so sz ≈ 3.728 a.

- **PML**: mp.PML(dpml, direction=mp.Z) — absorbing boundary **only along z** (the propagation direction). No PML in x or y.

- **Lateral boundary condition**: k_point = mp.Vector3() (i.e. k = 0). With no PML in x/y, this makes the simulation **Bloch-periodic in x and y at normal incidence** — MEEP tiles the single 528 nm × 528 nm disordered block **periodically to infinity** in the lateral directions. **This is not a finite, isolated disordered sample**; it is an infinite periodic array of one particular finite random tile. For a quasi-ordered but genuinely aperiodic biological structure, periodic tiling of a small tile (here ≈2.9 periods per side) can introduce spurious long-range order and artificial coherence at the tile boundary that would not be present in a real, non-periodic feather cross-section or in an ensemble/finite-domain experiment. We flag this explicitly as a modelling approximation, not a property of the real material.

- **Source**: mp.GaussianSource(fcen, fwidth=df), component=mp.Ex, a normal-incidence plane wave spanning the full lateral cell (size=mp.Vector3(sx, sy, 0)), placed at src_z = -0.5*sz + dpml + 0.2.

- **Frequency range**: λ = 0.36–0.78 µm (360–780 nm), fcen/df set from fmin=1/0.78, fmax=1/0.36; nfreq = 100 flux points.

- **Material**: keratin = mp.Medium(index=1.56), air = mp.air (n = 1.0); the voxel volume is supplied to mp.MaterialGrid(..., weights=design) (design = flattened 0/1 voxel array). do_averaging is left at the MEEP default (False, i.e. no subpixel smoothing) unless --subpixel-smoothing is passed.

- **Two-step reflectance recipe**: (1) an empty (default_material=mp.air) normalisation run records the incident flux and flux data via sim.get_flux_data; (2) sim.reset_meep(), then a second run with the structure geometry uses sim.load_minus_flux_data to obtain the reflected flux; R(λ) = -reflected/incident.

- **Termination**: mp.stop_when_fields_decayed(20, mp.Ex, (0,0,refl_z), 1e-3) in both runs.

#### S5.4 Exact commands

Baseline run reported in the main text (Methods §2.8 / Results §3.5), resolution = 24, seed = 1:

cd model

python fdtd3d_meep.py --d 185 --vox 24 --nm-per-vox 22 --res 24

Resolution-sweep commands (same d/vox/nm-per-vox, --res varied):

python fdtd3d_meep.py --d 185 --vox 24 --nm-per-vox 22 --res 24

python fdtd3d_meep.py --d 185 --vox 24 --nm-per-vox 22 --res 36

python fdtd3d_meep.py --d 185 --vox 24 --nm-per-vox 22 --res 50

Subpixel-smoothing commands (--subpixel-smoothing) for Table 2 are:

python fdtd3d_meep.py --d 185 --vox 24 --nm-per-vox 22 --res 24 --seed 1 --subpixel-smoothing

python fdtd3d_meep.py --d 185 --vox 24 --nm-per-vox 22 --res 36 --seed 1 --subpixel-smoothing

python fdtd3d_meep.py --d 185 --vox 24 --nm-per-vox 22 --res 50 --seed 1 --subpixel-smoothing

The Table 3 seed sweep uses the same command with --seed 1, --seed 2, or --seed 3, and --d 140 or --d 250, at --res 24 --subpixel-smoothing. All other parameters (cell, PML, source, frequency range, termination) are unchanged from §S5.3.

#### S5.5 Results

**Table 1 — resolution sweep, no subpixel smoothing (as committed; d = 185 nm, pore fraction 0.5, disorder 0.12, vox = 24, 22 nm/voxel, seed = 1).** Analytic 3-D structure-factor prediction for the same nominal parameters (structure3d.colour_3d, size = 48 for finer sampling): **540 nm**.

| Resolution (px/µm) | Peak wavelength |
|---|---|
| 24 | 477 nm |
| 36 | 381 nm |
| 50 | 576 nm |

**Not converged**: the peak varies by 195 nm (381–576 nm) across this resolution range with no monotonic trend.

**Table 2 — resolution sweep, with subpixel smoothing (do_averaging=True; same parameters as Table 1).**

| Resolution (px/µm) | Peak wavelength |
|---|---|
| 24 | 576 nm |
| 36 | 581 nm |
| 50 | 473 nm |

Smoothing changes the individual values substantially relative to Table 1 and is **also not converged** (108 nm spread, non-monotonic).

**Table 3 — seed sensitivity at fixed resolution (24 px/µm, do_averaging=True), individual results and summary statistics, for a blue-producing and a red-producing spacing.**

| d (nm) | seed 1 | seed 2 | seed 3 | mean | SD |
|---|---|---|---|---|---|
| 140 | 401 nm | 360 nm | 362 nm | 374.3 nm | 19.1 nm |
| 250 | 608 nm | 670 nm | 663 nm | 647.0 nm | 27.6 nm |

n = 3 seeds per spacing; not sufficient to establish a statistically robust trend, but consistent with larger dispersion at the longer (redder) spacing, The two-dimensional realisation analysis is reported separately in main text §3.4, Figure 6B; its dispersion does not increase monotonically with spacing.

#### S5.6 Interpretation and what would be needed for a quantitative claim

Across Tables 1–2, six independent (resolution × smoothing) settings for the *same* nominal geometry (d = 185 nm) give peaks ranging from 381 to 581 nm — a span that covers most of the blue-to-orange visible range. This is not resolution convergence in any usable sense, and, as currently run, the FDTD results cannot support a claim of "independent quantitative (or even independent qualitative) verification" against the analytic structure-factor prediction or against real plumage colour.

Causes we can identify from the data above:

1. **Small simulation volume.** 24 voxels at 22 nm/voxel = 528 nm per side is only ≈2.9 periods of a 185 nm structure; coherent scattering from disordered media is known to show large sample-to-sample (mesoscopic / speckle) fluctuations at this size, before any numerical-resolution issue is considered.

2. **Periodic (not finite) lateral boundary** (§S5.3): the simulation is a perfectly periodic tiling of one small random tile, not a finite or ensemble-averaged random medium — a structural mismatch with the physical system being modelled, independent of resolution.

3. **Under-resolution of the 22 nm voxel grid.** At resolution 24 px/µm the MEEP pixel size (≈41.7 nm) is coarser than the material's own 22 nm voxel spacing; subpixel smoothing (Table 2) changes the answer but does not fix convergence, consistent with the dominant source of variability being (1)-(2) rather than smoothing alone.

**What full convergence would require:** resolution swept well past the point where results stop changing (likely ≥100 px/µm here, at much higher compute cost), a lateral cell several periods larger with either absorbing (not periodic) lateral boundaries or explicit ensemble averaging over many periodic tiles, and averaging over many disorder realisations at each resolution (as begun in Table 3, but with far more than 3 seeds). This is identified as future work; it has not been done here.
